# Asynchronous firing and off-states in working memory maintenance

**DOI:** 10.1101/2025.10.12.681904

**Authors:** Rana Mozumder, Zhengyang Wang, Wenhao Dang, Junda Zhu, Benjamin M. Hammond, Anna Machado, Christos Constantinidis

## Abstract

Persistent spiking activity and activity-silent mechanisms have been proposed as neural correlates of working memory. To determine their relative contribution, we recorded neural activity from the lateral prefrontal and posterior parietal cortex of two male macaques using high-density microelectrode probes. We found that, when averaged across all neurons, persistent delay activity was observable throughout the duration of single trials in populations of prefrontal neurons with silent periods that did not deviate significantly from chance. However, temporal fluctuations in activity-dependent mnemonic information were present and weakly correlated between the prefrontal and posterior parietal cortices, suggesting at least partial, long-distance synchronization of off-states. Decoding accuracy of neurons recorded simultaneously was also reduced relatively to pseudo-populations constructed by splicing different trials together. Our results support an asynchronous state of working memory, maintained by the distributed pattern of persistent discharges across cortical neurons, which is subject to widely distributed fluctuations in information representation fidelity.

## INTRODUCTION

The neural basis of working memory, the ability to maintain and manipulate information over a period of a few seconds, has been debated (Constantinidis et al., 2018; Miller et al., 2018). Classic neurophysiological experiments identified neurons in the prefrontal cortex (PFC) and other brain areas that generate persistent discharges after stimuli are no longer present (Funahashi et al., 1989; Fuster and Alexander, 1971). The assumption that this persistent activity represents the neural correlate of working memory has been challenged, since (Lara and Wallis, 2014; Lundqvist et al., 2018). In particular, it has been pointed out that firing of individual neurons rarely spans the entire delay period of working memory tasks and the appearance of persistence may be an “artifact of averaging” (Lundqvist *et al*., 2018). Alternative mechanisms have been proposed, instead, that are based on short-term synaptic changes (Mongillo et al., 2008; Stokes, 2015), or the rhythmicity of neuronal discharges (Lundqvist et al., 2016), leading to the proposal that mnemonic information might be stored exclusively in synaptic weights, during transients when neuronal activity is silent (Panichello et al., 2024). Newer versions of the persistent-activity model counter that individual neurons may not be continuously active, but discharges in the network may still span the entire delay period of a working memory task, on a single-trial basis (Jaffe and Constantinidis, 2021). This asynchronous-state mode, inspired by theoretical models (Brunel, 2000; Hansel and Mato, 2003; Hansel and Sompolinsky, 1996; Renart et al., 2010; Tan et al., 2014), has only been speculated for persistent activity and has not been verified experimentally in the prefrontal cortex. In the absence of definitive evidence, the debate remains unresolved (Lansner et al., 2023).

Part of the challenge in pinpointing mechanisms of working memory maintenance has been the piecemeal approach of neurophysiological recordings from the prefrontal cortex, involving one or a few neurons at a time. Modern neurophysiological techniques have altered the landscape of experimental neurophysiology, making it possible to monitor the activity of hundreds of neurons at a time, e.g. with multi-contact probes known as Neuropixels (Jun et al., 2017). Such advances now make it possible to test experimentally long-debated ideas about persistent activity – as well as understand the dynamics of neuronal activity across populations (Ebitz and Hayden, 2021). We were therefore motivated to obtain large-scale recordings from the prefrontal cortex of monkeys performing working memory tasks. We examined population activity in individual trials across simultaneously recorded neurons. We were thus in a position to test the role of neural activity during working memory maintenance across different cortical sites.

## RESULTS

We recorded neural activity from the prefrontal cortex of two monkeys trained to perform the Oculomotor Delayed Response task (ODR – Fig. 1A). We analyzed results from 13 recording sessions (8 from monkey P and 5 from monkey V) sampled from the dorsolateral prefrontal cortex of the two animals (Fig. 1B). The animals achieved performance of approximately 90% correct trials during these sessions (Fig. 1C). In an initial series of recordings data were obtained from the dorsolateral prefrontal cortex with Neuropixel electrodes (Fig. 1D-left), and more than 80 single units were identified from each session (Fig. 1D-right). A total of 2297 single units were available for analysis (mean: 176.7 per penetration, standard deviation: 83.5). In addition, 837 multi-units were available (mean: 64.4 per penetration, standard deviation: 27.8).

**Figure 1.**
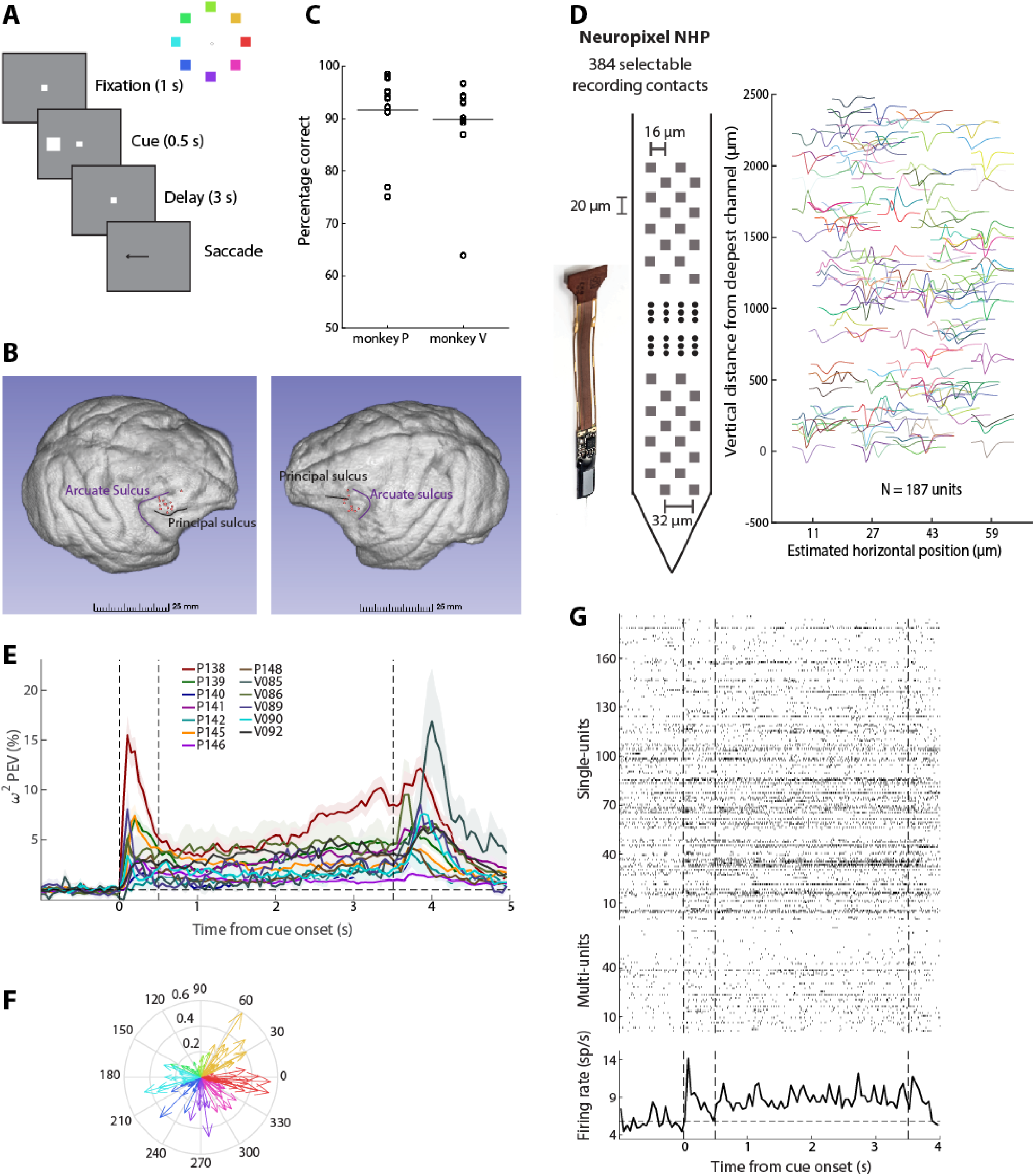
Task and recordings. (A) Left: Sequence of events in the ODR task. The monkey is required to maintain fixation for 1 s to observe the cue stimulus, remain fixated for the delay period of 3 s, and make a saccade to the remembered cue location after the delay period, indicated by the disappearance of the fixation point. The monkeys were rewarded for the successful completion of the trial. Right: Cue stimuli are shown in a ring with different colors. (B) Recording locations (red dots) are shown on the surface of the brains of monkey V (left) and P (right). (C) Percentage of correct trials for each recorded session (black circles). The horizontal lines indicate the mean across sessions (91.64% and 89.82% for monkey P and V respectively). (D) Left: Schematic of Neuropixel NHP probe. Right: Recorded spike waveforms of 187 single units extracted from an example session from monkey P. (E) Mean percentage of explained variance (PEV) across neurons selective for the delay period extracted from individual recording sessions. Shaded areas indicate the mean ± standard error of the mean (s.e.m.). Dotted vertical lines indicate stimulus onset (0 s), delay period onset (0.5 s), and end of delay period (3.5 s) respectively. (F) Compass plot showing tuning of all single units selective for the delay period recorded in an example session (P138). Each arrow’s direction denotes the angle of tuning and length indicates the strength of tuning for each unit. Each arrow color indicates the spatial selectivity of that neuron. (G) Top: Raster plot of single units for an example trial (315° cue stimulus as shown by the ring on the left) recorded simultaneously for the same recording session as in F. Each row shows the spiking timestamps of individual units. Middle: Same convention as the top panel for multi-units from the same example trial. Bottom: Peri-Stimulus Time Histogram (PSTH) for the same example trial. Dotted horizontal line shows the average baseline activity. Dotted vertical lines showing stimulus onset (0 s), delay onset (0.5 s), and end of delay period (3.5 s) respectively.

### Individual sites exhibit selective persistent activity

Firing rate and spatial selectivity properties of neurons obtained from such laminar recordings were generally consistent with previous studies that sampled neurons sequentially. Selectivity in the task varied considerably across sessions, quantified with the mean, bias-corrected percentage of explained variance (ω^2^ in Fig. 1E). However, across all sessions, an above 0 mean value was evident throughout the entire delay period of the task. Averaged across the entire delay period, in all 13 sessions, the mean ω^2-^PEV value was significantly greater than 0 (one-sample *t*-test, *p*<0.03 in every case). Tuning of individual neurons was also highly variable (Fig. 1F), rather than representing the same spatial location, consistent with previous studies in the prefrontal cortex revealing no strong topographic organization of visual space across sites (Constantinidis et al., 2001a; Mendoza-Halliday and Martinez-Trujillo, 2017). Nonetheless, individual sites exhibited an overall bias towards different locations, and there was an overweighting of contralateral locations. Preferred locations of all neurons across all sessions are shown in Fig. S1. Neurons with selective persistent activity were observed across the depths of the cortex, although slightly higher levels of mean activity at the end of the delay period were present in superficial layers (Fig. S2), in agreement with prior studies (Markowitz et al., 2015; Zhu et al., 2023).

We tested directly the existence of the asynchronous state of persistent activity maintenance by quantifying population firing rate, pooled across all neurons recorded in single trials. An example is shown in Fig. 1G for a stimulus presented at the 315° (bottom right) location. Although persistent activity was barely evident in single units, across the population of 253 units (187 single units and 66 multi-units) robust persistent activity was present throughout the entire trial. Population neuronal firing rate was elevated throughout the delay period compared to the fixation period (for 57/60 50-ms bins during the delay period firing rate exceeded the threshold of significance; paired t-test, False Discovery Rate (FDR)-corrected at the q=0.05 level). Additional examples of single trials with persistent activity are shown in Fig. S3. It is critical to point out that robust persistent activity was not present for locations away from the overall preferred location of the site. Had we not identified the overall preferred location for this site and plotted responses to a random location, our conclusion might have been absence of persistent activity. Indeed, an example of non-preferred location from the same site is shown in Fig. S3B which did not show persistently elevated delay activity (only 1 time bin exceeded the significance threshold, paired t-test, FDR-corrected at the q=0.05 level). We have pointed out the logical fallacy of drawing a negative conclusion about the presence of persistent activity when using a limited stimulus set, precisely for this reason (Constantinidis *et al*., 2018).

The presence of elevated activity during the trial does not necessarily imply that this activity reliably encodes the stimulus information. We therefore performed decoding analysis to determine how reliably the stimulus information could be decoded from the neuronal firing. A classifier was trained and tested separately at each time point to decode a stimulus from all other stimuli (Fig. 2). On average, robust decoding was present when we trained and tested on the same time points (Fig. 2-central panel), though the absolute performance differed between stimulus conditions, depending on the overall preference of the site (contrast Fig. 2-lower right and upper left panels; each peripheral panel indicates decoding results of trials with the corresponding cue from all other trials). Averaged decoding results for all sessions are shown in Fig. S4.

**Figure 2.**
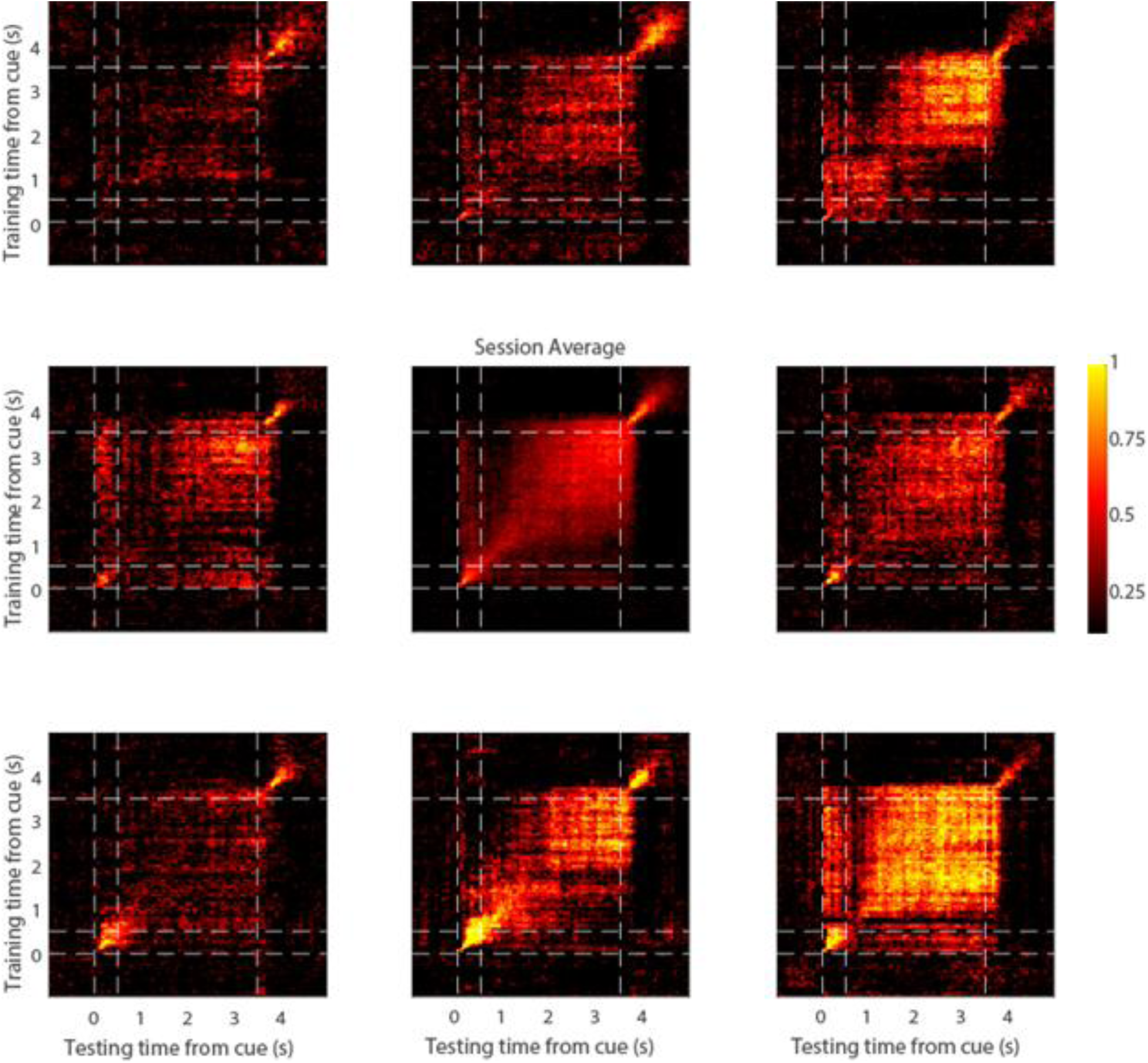
Cross-temporal decoding. Decoding performance is shown from a single session (P138). Eight peripheral panels correspond to the eight cue locations. Each plot shows the average decoding performance for trials of that stimulus. The vertical and horizontal dotted lines represent stimulus onset (0 s), delay onset (0.5 s), and end of delay period (3.5 s) respectively. Center panel represents the mean decoding performance across all trials of the session.

### Inter-Spike Interval analysis

We also sought to test whether prolonged silences occur within individual trials, during which persistent activity ceases. To directly test for the presence of such population-level silent periods, we compared the maximum inter-spike intervals (maxISIs) from the empirical data to a shuffled null distribution (see Methods). The central hypothesis is that coordinated silences across neurons would disrupt the typical inverse relationship between firing rate and maxISI (Fig. 3A-D). Specifically, in the presence of long enough silent periods, log₁₀(maxISI) would be dominated by the length of the silent periods and be less dependent on log₁₀(firing rate), resulting in a flatter empirical slope than expected by chance (Fig. 3E-H). In contrast, trial-shuffling breaks population-level coordination and restores a more typical negative linear relationship between the logarithm of maxISI and the logarithm of firing rate (blue line in Fig. 3H).

**Figure 3.**
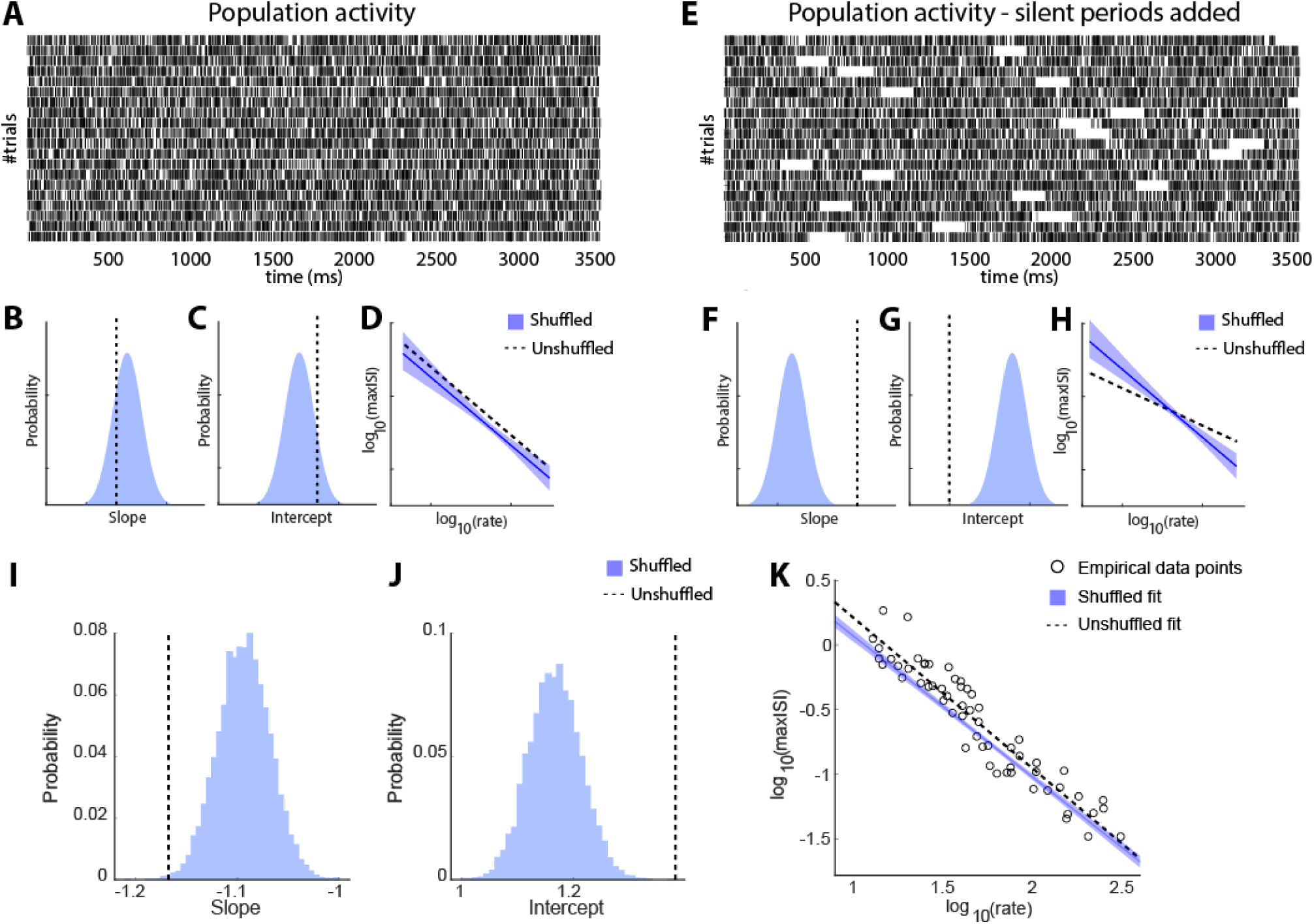
Inter-Spike Interval analysis. (A) Raster plot for simulated, population data, with no silent periods. For each trial, the activity of all simulated neurons was pooled together to create the population activity. (B-C) Distribution of slopes (B) and intercepts (C) of log-transformed maximum ISIs vs. log-transformed firing rate, for the shuffled populations with no silent periods. Dotted vertical lines indicate the slope and intercepts for the unshuffled populations. (D) Logarithm of the maximum ISI is plotted against the logarithm of population firing rate in the population. Blue solid and black dotted line indicate regression lines for shuffled and unshuffled data. (E) Raster plot for population data as in A, but for a population with silent periods added. (F-H). Same as in B-D, for populations with silent periods. (I-J) Same as in B and C, but for the empirical data constructed from populations of at least 5 neurons. K) Logarithm of max-ISIs is plotted against the logarithm of firing rate. The blue solid line and black dotted line indicate the lines fitted to the shuffled and unshuffled data. The shaded region denotes the 95% confidence interval, and the circles indicate the data points for the empirical data line.

To validate this approach, we generated a simulated dataset consisting of two types of populations: one without silent periods and one in which silent periods were imposed randomly. Example rasters for both conditions are shown in Fig. 3A, E. Populations without silent periods showed slopes and intercepts closely matching the shuffled distribution (schematically shown in Fig. 3B-C), resulting in overlapping log-log relationships (Fig. 3D). In contrast, populations with silent periods exhibited shallower empirical slopes and lower intercepts relative to the shuffled distribution (Fig. 3F-G), producing a flattened log₁₀(maxISI) – log₁₀(firing rate) relationship (Fig. 3H).

With these expectations, we employed the same analysis on our recorded dataset, examining the inter-spike intervals of populations of neurons consisting of at least 5 delay-period selective neurons that shared the same preferred stimulus location (Fig. 3I-K). We saw no evidence of a steeper slope for the shuffled distribution that would have been indicative of population silent periods. In fact, the empirical data exhibited a steeper slope compared to the shuffled data (one-sided permutation test; p=0.002), opposite in sign to what is expected of the off-period hypothesis and also different from that expected of independent, Poisson-like spiking. This result implies that the population spiking is more regular than a Poisson-like mode, and instead of turning on and off synchronously, the neurons encoding for the same cue information are coordinated to tile the delay period. Qualitatively similar results were obtained when we varied the size of populations examined to obtain the population ISIs (n≥2 to n≥10, Fig. S5A-B). We similarly examined ISIs in error trials (Fig. S5C-E). The slopes of the empirical distributions tended to be slightly higher than those of correct trials, implying that trials contained longer ISIs tended to result in errors, but even in this case we found no evidence of coordinated silent periods of spiking; the slope of permuted data was not steeper than the empirical data (one-sided permutation test; p=0.12 for n≥5, in Fig. S5D).

### Coordinated off states between areas

The analysis presented so far relied on raw firing rate, which provides only a rough estimate of information represented by the neuronal population. For example, in Figures 1 and 3, temporary spiking of neurons whose preferred location differs from the actual cue location may contribute to the apparent absence of absolute gaps in firing rate but it does not follow that information about the cue location is maintained precisely. In fact, evidence exists that mnemonic information decodable from neuronal activity fluctuates, resulting in alternating transients of “on” and “off” state in populations of neurons recorded by laminar probes (Panichello *et al*., 2024). One caveat in assuming these transient fluctuations to be true lapses in activity-dependent memory code is that activity of neurons sampled by a laminar probe, roughly corresponding to a column in the prefrontal cortex, likely exhibit stronger correlations compared to the activity of neurons located further apart on the cortical surface, while it is well understood that multiple areas participate in the maintenance of working memory and reverberation across long-range connections is essential for activity to enter a multi-stability regime for persistent activity (Mejias and Wang, 2022). It was important, therefore, to test for the existence of coordinated on and off-states across areas. For this purpose, we obtained an additional sample of neurons recorded simultaneously from the PFC and the posterior parietal cortex (PPC). We recorded a total of 810 units in the PFC and 836 units in the PPC across 7 sessions, sampling both regions simultaneously. The PFC recordings consisted of 621 single units and 189 multi-units, while the PPC recordings included 501 single units and 335 multi-units, all from the same hemisphere. Neurons in both areas had a wide range of preferred locations (Fig. S6).

To identify transient fluctuations in mnemonic information, we followed a binary decoding approach where we classified each cue location from its diametric location using the population of neurons recorded within each region respectively (Fig. 4A; and Fig. S7). For each trial, we labeled segments in the delay period as either on- or off-states depending on whether classifier confidence was high or low compared to a null distribution (Fig. 4B - see methods). Off-states defined in this fashion, were represented by a flattening of neuronal tuning curves, resulting in an average 77.6% reduction in tuning depth (difference between peak and baseline normalized firing rate, Fig. 4C). This led to the on-state firing rate (red trace in Fig. 4D) being significantly greater than that of the off-state (blue trace) firing rate (paired t-test: *t*_633_=9.1, *p*=1.21E-18), as well as the mean firing rate (black trace) compiled from all trials (paired t-test: *t*_633_=10.566, *p*=3.84E-24). Similarly, we found a reduction of 75.7% in tuning depth in PPC (Fig. 4G) and corresponding differences in firing rates (paired t-test: *t*_707_=10.839, *p*=1.984E-25 for on- vs. off-state; *t*_707_=11.998, *p*=2.552E-30, for on vs. mean).

**Figure 4.**
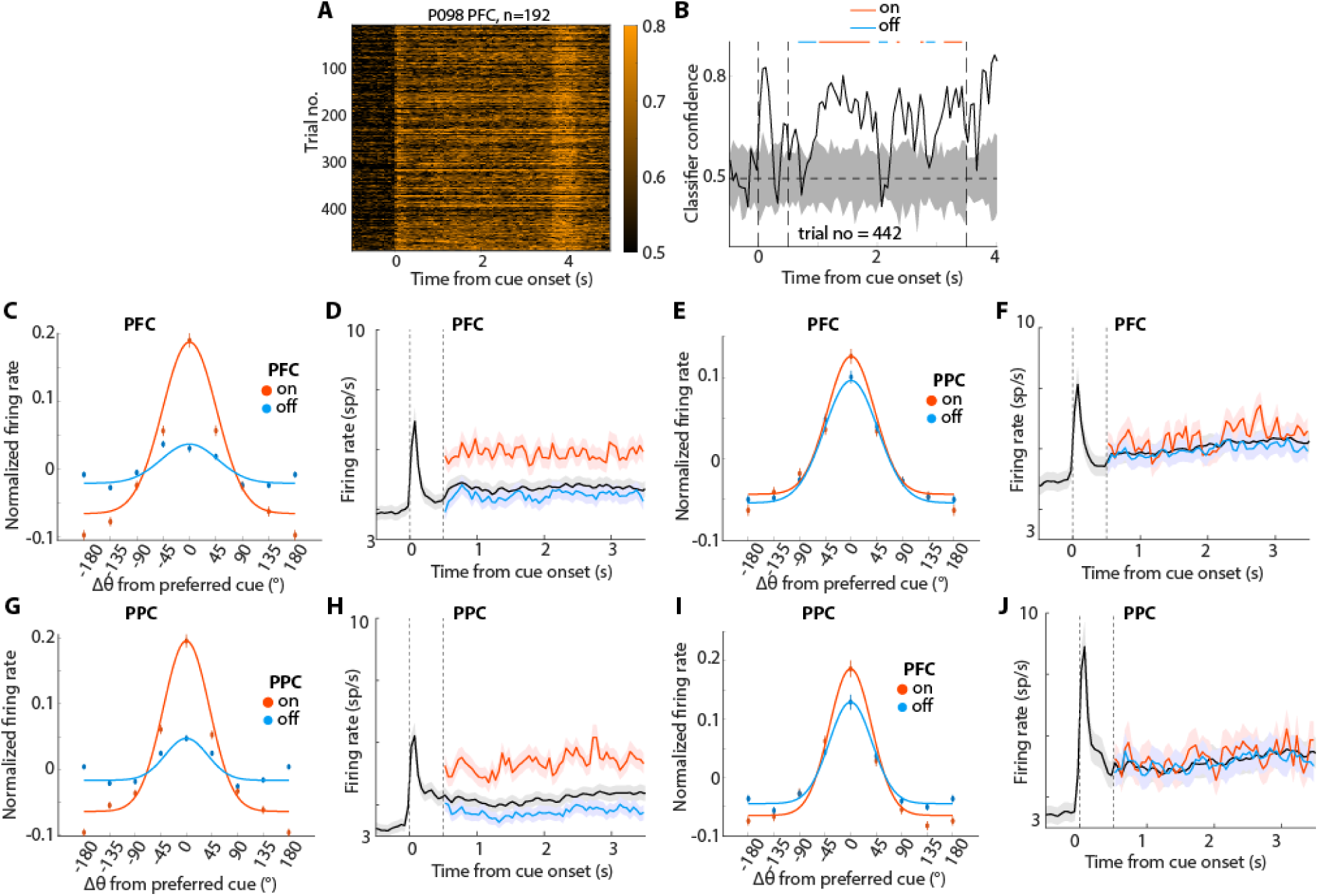
Changes in tuning properties and firing rate during On and Off states. (A) Single-trial classifier confidence for all trials of an example session from simultaneous PFC-PPC recordings. (B) Labeling of on and off states on the confidence time series of an example trial from A. Gray shaded area shows the shuffled distribution. Dotted vertical lines show stimulus onset (0 s) and delay onset (0.5 s). The dotted horizontal line indicates chance level (0.5). (C) Tuning functions (firing rate z-scored across trials) for on and off states for PFC neurons. (D) Mean population firing rates for the preferred location of PFC neurons as in C. Dotted vertical line indicating onset of and disappearance of cue respectively. (E-F) Tuning functions and mean population firing rate of PFC activity determined based on the on- and off-states of the PPC recording. (G-H) Similar to C-D but for PPC recordings. (I-J) Tuning functions and mean population firing rate of PPC activity determined based on the on and off-states of the PFC recording.

When we examined PFC firing in periods of off-states defined by PPC activity, we also found a decrease in tuning depth (11.7%, Fig. 4E), which was significant across neurons (one-sample t-test, t_489_=3.24, p=0.001). We similarly detected a significant decrease in PPC tuning depth during off-states defined by PFC activity (mean: 30.3%, Fig. 4I, t_378_=5.45, p=9.1E-8). On and off states detected in one region were characterized by only subtle differences in firing rate in the other (Fig. 4F, J). Still, a PFC firing rate difference was present between on and off states defined by PPC activity (paired t-test: *t*_489_=3.52, *p*=4.75E-4) or between on and mean rates (*t*_489_=3.757, *p*=1.93E-4), though PPC activity based on PFC on- vs. off-state comparison did not reach statistical significance (paired t-test: *t*_378_=0.67, *p*=0.504 for on- vs. off-state comparison; *t*_378_=0.93, p=0.354 for on- vs. mean rate comparison).

We further tested whether on- and off-states represent phases of oscillatory firing rate, but no obvious rhythmicity was present in their incidence (Fig. S8). In terms of their influence on behavior, off-states exhibited a weak negative relationship with reaction time, which only reached statistical significance for the prefrontal cortex (Fig. S9, one-sided bootstrap test, p=0.033 for PFC; p=0.136 for PPC). The relative number of on- and off-states during the delay period also varied between correct and error trials and more so for the PFC than the PPC (Fig. S9B-C). In correct trials, on-states were more frequent when the stimulus appeared at the overall preferred location of the site (location for which highest decoding confidence was achieved), compared to the non-preferred location (lowest confidence); in error trials, the opposite was true (red boxes in Fig. S9B). These relationships were reversed for off-states (blue boxes in Fig. S9B). A 3-way ANOVA revealed a significant interaction between correct/incorrect status and preferred/non-preferred location (F1,32=7.74; p=0.009); and between preferred/non-preferred location and on/off state (F1,32-9.83, p=0.004); as well as a 3-way interaction (F1,32=19.5; p=1.1E-4). A similar overall pattern was present in the PPC (Fig. S9C), however none of these interactions reached statistical significance (p>0.1 in each case).

These results suggest that although off states are not readily evident in firing rate measures pooling activity across neurons and locations, temporal fluctuations in activity-dependent mnemonic information represent a global and coordinated feature, likely reflecting the dynamic nature of brain states, and exert some influence on behavior.

### Simultaneous vs. pseudo-population activity

To explore further the implications of off-states in neuronal encoding, we examined populations of neurons recorded simultaneously and asynchronously. In principle, single-unit spike timing may exhibit features of irregularity or stochastic oscillations (Compte et al., 2003; Pesaran et al., 2002), but the network as a whole may still maintain an attractor state (Compte et al., 2000; Lundqvist et al., 2010). On the other extreme, coordinated periods of off-states may interrupt irretrievably information maintenance in the network. Under this scenario, splicing neurons recorded in different trials would inflate decodable information due to the averaging out of non-matching fluctuations in different trials, in essence, removing the true silent periods of individual trials and artifactually creating an illusion of uninterrupted activity (Lundqvist *et al*., 2018). To distinguish between the two alternatives, we compared the decoding accuracy of neurons recorded simultaneously in a session to that of 100 pseudo-populations constructed by shuffling neuronal activity across trials within each stimulus condition, thereby preserving stimulus identity while disrupting within-trial correlations between neurons (Fig. 5).

**Figure 5.**
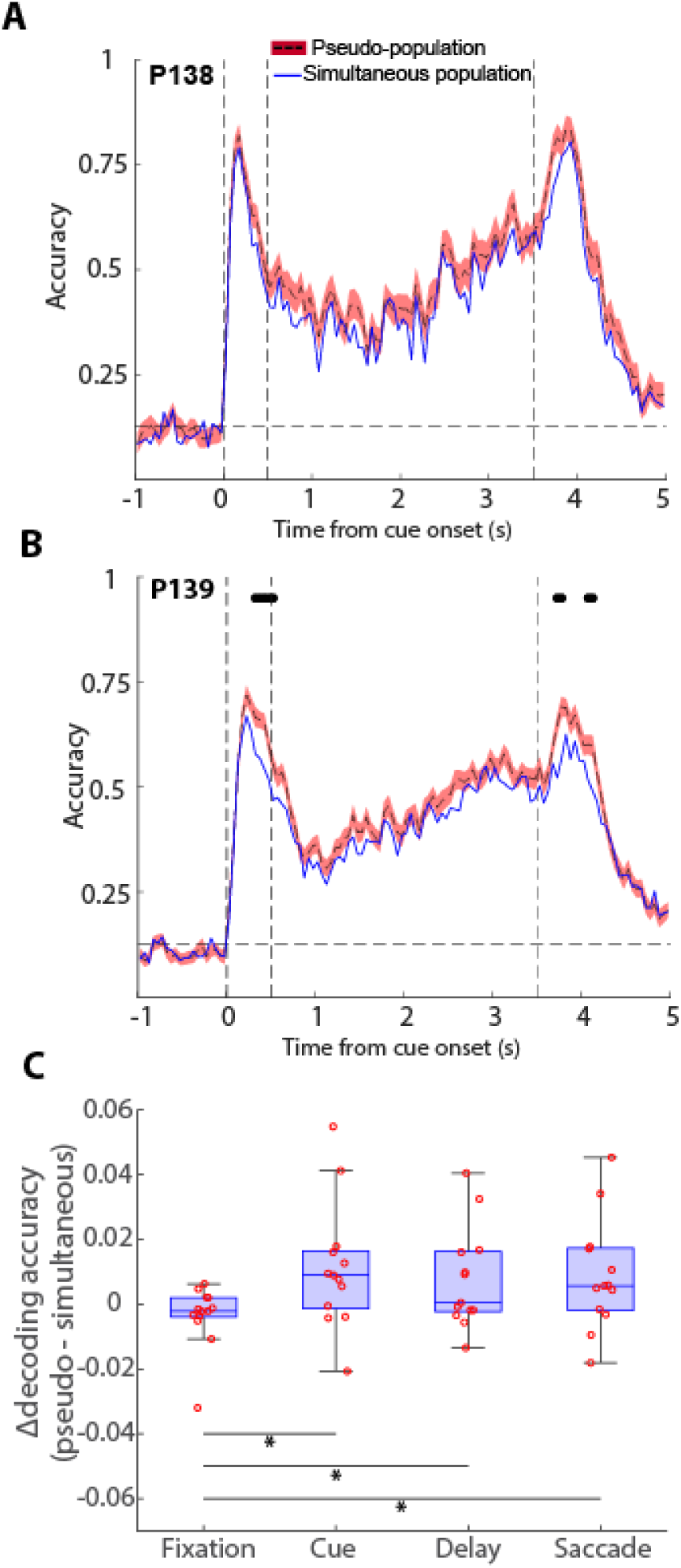
Pseudo-populations vs. simultaneous decoding. (A-B) Two example sessions were obtained using decoding the simultaneous recordings compared to 100 pseudo-populations where neuronal activity was randomly shuffled across trials. Black circles at the top of each subplot denote significant difference in simultaneous and pseudo-population decoding. Multiple comparisons across time points were corrected using the Bonferroni method to control for Type I error. The vertical dashed lines represent stimulus onset (0 s), delay onset (0.5 s), and end of delay period (3.5s) respectively. The horizontal dashed line denotes chance-level decoding accuracy. C. Mean difference in decoding accuracy for each of the task epochs. Points represent individual sessions.

Two example PFC sessions comparing the simultaneous and pseudo-population decoding are shown in Fig. 5A and B. In support of the idea that asynchronous firing rate reasonably approximates the synchronous one, the absolute value of decoding in the two populations was very similar. However, the simultaneous population was consistently lower than the pseudo-population, a difference that was highly significant (Wilcoxon signed-rank test, p=1.6E-17, Fig. 5A). Across all sessions examined (Fig. 5 and Fig. S10), the decoding performance of the simultaneous population was significantly lower in 7/13, significantly higher in 1/13, and not significantly different in 5/13. Importantly, the relative decrease in decoding accuracy in the simultaneous populations was not limited to the delay period alone. A similar decrease was present during the cue presentation, when a physical stimulus was present in the screen, and during the response epoch when a saccade was performed, rather than only in the delay period when maintenance of information depended entirely on activity reverberating in the network, and this was also true for individual timepoints where information significantly differed from each other were rare (black dots in Fig. 5B, and Fig. S10). Decoding accuracy averaged across the entirety of each of the task epochs revealed that the overall difference in decoding accuracy between the real population and pseudo-population was not significantly different for the delay period compared to the cue and saccade periods (Fig. 5C). The result indicates that coordinated periods of decreased neural information, e.g. due to variations in neuromodulatory tone (Nassar et al., 2012), are present during normal firing, but do not lead to intervals of absolute cessation of delay-period activity.

## DISCUSSION

Our results demonstrate that asynchronous firing in a population of cortical neurons is sufficient to maintain information about a remembered stimulus. Persistent firing, elevated above the baseline firing rate during the fixation period, was evident by raw pooling of activity across all neurons sampled by a multi-contact probe, even considering that some neurons are expected to be suppressed during the delay period of working memory tasks (Constantinidis and Goldman-Rakic, 2002; Zhou et al., 2012). The choice of stimulus for the demonstration of persistent activity was critical for each recording site. Non-optimal stimuli may result in ostensible lack of persistent activity. However, selecting the optimal stimulus from eight equidistant spatial stimuli across all quadrants of the visual field was sufficient to elicit above-chance persistent activity for every site sampled in the dorsolateral prefrontal cortex. Despite the asynchronous state of firing, some off-states were evident in neuronal encoding, in agreement with prior studies (Panichello *et al*., 2024), which we now show they are coordinated between areas, and were associated with a higher likelihood of an error. Firing pauses during off-states were not absolute, however, and the activity across the distributed network maintained information about the stimulus being remembered.

### Competing models of working memory

Working memory is a core cognitive process that enables not only the storage of information over a timescale of seconds but also the execution of mental functions that require manipulation of information and contemplation of alternatives (Baddeley, 2012). Originally thought to reside within the prefrontal cortex, it is now understood to engage a broader network of brain areas, including the sensory pathways connected to it (Jaffe and Constantinidis, 2021). However, prefrontal cortex remains central for working memory function and neurons manifesting neural correlates of working memory function are readily encountered in this area.

Several network architectures can give rise to persistent activity (Chaudhuri and Fiete, 2016; Zylberberg and Strowbridge, 2017), though simple networks with local recurrent excitation and broader feedback inhibition have been shown to function as “bump attractors” and generate persistent discharges that capture many empirical properties of prefrontal neurons (Compte *et al*., 2000). Experimental evidence suggests that persistent discharges are causally related to behavior. Trials in which persistent activity is diminished are more likely to result in errors (Funahashi *et al*., 1989; Zhou et al., 2013). A monotonic relationship exists between behavioral performance and persistent activity in tasks that parametrically modulate the difficulty of a working memory judgment (Constantinidis et al., 2001b). Choice probability analysis, comparing the distributions of firing rates in the delay period of correct and error trials, also reveals a strong relationship between prefrontal persistent activity and the behavioral outcome of each trial (Mendoza-Halliday et al., 2014). Trial-to-trial deviations in memory-guided eye movements can be traced to differences in neural activity among prefrontal neurons that generate persistent activity (Wimmer et al., 2014).

Some neurophysiological experiments have reported that individual neurons are only transiently representing information in the delay period (Zaksas and Pasternak, 2006) and persistent activity can be highly variable during the course of a trial, and from trial to trial (Lundqvist *et al*., 2016). This led to the idea that previous reports of persistent activity were an artifact of averaging. Studies involving dual tasks or retro-cue paradigms that require subjects to maintain actively one stimulus for some period of time in a trial have shown that neural activity returns to baseline when a stimulus is not actively maintained, but may increase after the subject is cued, suggesting that information may be latent and yet have the ability to be reactivated (Rose et al., 2016; Watanabe and Funahashi, 2007; Wolff et al., 2017). Such findings raised the possibility that working memory can be maintained in an ‘activity-silent’ form. Mechanistic models have been proposed to account for such experimental findings. These included models relying on dynamic encoding so that the precise pattern of activation of an ensemble of neurons at each time point during a working memory task can be used to decode the identity of the stimulus even though overall activity during the delay period is not significantly elevated above the baseline (Stokes et al., 2013). Another class of “activity-silent” models rely on synaptic mechanisms. Activity during encoding temporarily changes synaptic efficacy within the neural network, leaving behind a temporary cellular memory trace (Mongillo *et al*., 2008; Sugase-Miyamoto et al., 2008). Such mechanisms may be mediated by calcium availability at the presynaptic terminal, whose kinetics have a time constant in the scale of seconds, or other synaptic factors (Mongillo *et al*., 2008). While such activity silent mechanisms may support working memory maintenance, it has been argued that more complex processes, such as manipulating information in working memory require spiking activity as intracellular signals such as Calcium concentration are trapped in synapses and must be reactivated by other incoming spikes (Wang, 2021).

### Asynchronous firing and off-states

The asynchronous state of neural firing, characterized by high variability, and independence of firing between nearby neurons, was initially characterized in the visual cortex (Hansel and Sompolinsky, 1996). Simulations of sparsely connected networks of excitatory and inhibitory neurons also confirmed that integrate-and-fire units alternate between synchronized and asynchronous states (Brunel, 2000; Hansel and Mato, 2003). Theoretical work demonstrated that recurrent neural networks could generate an asynchronous state characterized by arbitrarily low spiking correlations despite substantial amounts of shared input because fluctuations in the activity of excitatory and inhibitory populations balance each other finely (Renart *et al*., 2010). Networks with strongly connected units display rich internal dynamics, in which the firing rates of individual neurons fluctuate strongly in time and across neurons (Ostojic, 2014). Somewhat paradoxically, sensory stimulation has been shown to shift the firing of visual neurons to asynchronous states (Tan *et al*., 2014).

In the context of working memory, existence of an asynchronous state has long been speculated. It was argued on theoretical grounds that failure to detect persistent activity across every trial in individual neurons are consistent with computational models of persistent activity: Across the population of prefrontal neurons, only a small minority would be expected to be active during maintenance of any single stimulus in memory and variability in discharge rate during the course of the trial would be expected even among neurons that are highly active at some time point, as the activity might drift in the population (Jaffe and Constantinidis, 2021). Other, indirect, evidence have also suggested existence of an asynchronous state, e.g., by the increased variability of spiking in the delay interval following a non-preferred stimulus in the receptive field of a neuron (Compte *et al*., 2003).

The present results provide direct experimental validation of this view. We show that asynchronous firing accounts well for persistent activity across the population. The collective activity of a site remained elevated above baseline when the overall preferred stimulus was maintained in memory but the same sites were mostly silent for non-optimal stimuli. The common practice of estimating decoding accuracy based on pseudo-populations of neurons recorded asynchronously (Meyers et al., 2012) was found to be a reasonable assumption, in agreement with prior studies that have also decoded robust activity from simultaneously recorded neurons (Leavitt et al., 2017). Weak, positive correlations between neurons recorded simultaneously are well understood to limit the ability to extract information from ambiguous stimuli by pooling increasingly larger populations of neurons (Cohen and Kohn, 2011; Zohary et al., 1994). However, in the case of unambiguous stimuli such as those used in the present study, correlated firing that has described for the prefrontal cortex (Leavitt *et al*., 2017; Qi and Constantinidis, 2012) had minor effects, leading to similar decoding performance of simultaneously recorded populations and pseudo-populations of prefrontal neurons.

Although we have emphasized the importance of persistent discharges, we do not wish to imply that activity-silent and rhythmic mechanisms do not exist. A well-established example involves the serial bias between successive trials which is present in the ODR task, even though tuned persistent activity disappears in the inter-trial interval following the end of a trial (Barbosa et al., 2020). In our current study, we identified off-states, which were present within sites, and across cortical areas, possibly due to variations in neuromodulatory tone (Nassar *et al*., 2012). On- and off-states in our dataset did not have obvious rhythmicity to suggest that they coincide with phases of oscillatory firing. The existence of off-states suggests that short-term plasticity mechanisms may play a role in sustaining activation even when firing subsides, though less effectively than persistent discharges, as they were more likely to result in errors. Overall, our results suggest that synaptic mechanisms appear to influence behavior, but ultimately their influence may depend on their ability to modulate spiking output (Jaffe and Constantinidis, 2021).

### Limitations of the study

Linear probes allow recordings from large numbers of neurons, however, these are concentrated in specific sites and we were thus unable to sample disparate locations within each cortical area. For the same reason we cannot exclude the possibility of off-states being related to traveling waves, which have been described in the prefrontal cortex and other areas (Davis et al., 2020; Zanos et al., 2015). Additionally, our study focused exclusively on spatial working memory. Whether similar mechanisms of asynchronous firing and off-states operate for object working memory, or whether the latter is qualitatively different is still an open question (Jaffe and Constantinidis, 2021), to be addressed in future studies.

## RESOURCE AVAILABILITY

### Lead contact

Further information and requests for resources should be directed to and will be fulfilled by the lead contact, Christos Constantinidis

### Materials availability

This study did not generate new unique reagents.

### Data and code availability

Data and original code used to analyze the data in this paper will be made available prior to publication. Additional requests may be directed to the lead contact.

## STAR * METHODS

## KEY RESOURCES TABLE

| REAGENT OR RESOURCE | SOURCE | IDENTIFIER |
| --- | --- | --- |
| Deposited data |  |  |
| Figure Data | This paper | Pending |
| Experimental models: Organisms/strains |  |  |
| Rhesus macaques (*Macaca mulatta*) | Alpha Genesis | N/A |
| Software and algorithms |  |  |
| MATLAB | Mathworks | R2019a-2024b |
| Other |  |  |
| Neuropixel probes | Neuropixels | NP1015 and NP1010 |
| 128-channel probes | Diagnostics Biochips | DA128-2 |

## EXPERIMENTAL MODEL AND SUBJECT DETAILS

Two male, ∼7 years old rhesus monkeys (Macaca mulatta) weighing 9-10 kg were used in this study. Monkeys were single-housed in communal rooms, and they had sensory interactions with other monkeys. All surgical and animal use procedures were reviewed and approved by the Vanderbilt University Institutional Animal Care and Use Committee, in accordance with the U.S. Public Health Service Policy on Humane Care and Use of Laboratory Animals and the National Research Council’s Guide for the Care and Use of Laboratory Animals.

## METHOD DETAILS

### Experimental setup

The monkeys sat with their head fixed in a primate chair while viewing a monitor (Samsung QM32C, 1920×1080 resolution, 60 Hz) positioned 69 cm away from their eyes with dim ambient illumination and were required to fixate on a 0.1° white square (fixation point) appearing in the center of the screen. To receive a liquid reward (water), the animals had to maintain fixation on the square while visual stimuli were presented at peripheral locations. The monkeys needed to maintain fixation within a 3° window centered around the fixation point. Any break of fixation immediately terminated the trial, and no reward was given. Eye position was monitored throughout the trial with an infrared eye tracking system (ISCAN ETL-200; ISCAN, Inc., Woburn, MA). The system has a precision of 0.5° when tracking the pupil. Eye position was sampled at 500 Hz, digitized, and recorded. The visual stimulus display, eye position monitoring, and stimuli synchronization with neurophysiological data were performed with in-house software implemented in the MATLAB environment (MathWorks, Natick, MA), using the Psychophysics Toolbox (Meyer and Constantinidis, 2005).

### Behavioral task

The monkeys were trained to perform an oculomotor delayed response (ODR) task. The trials started with the monkeys fixating on the fixation point. The task required them to remember the location of a cue stimulus flashed on a screen for 0.5 s (Fig 1A). The cue stimulus was a 1° white square appearing at one of eight locations arranged on a circle of 10° eccentricity, spaced by 45°. After the delay period, the fixation point was extinguished, indicating the monkey to make a saccade to the remembered location of the cue within 0.6 s. The saccade needed to terminate on a 6° radius window centered on the stimulus, and the monkey was required to hold fixation within this window for 0.1 s. Animals were rewarded with water for the successful completion of a trial. For the simultaneous PFC and PPC recordings, a variant of the ODR task was used; instead of 8 equidistant stimuli, we presented 120 equidistant stimuli, each 3° apart from each other. For the analysis of this task, we grouped the stimuli into 8 groups to keep it consistent across tasks.

### Surgery and neurophysiology

A 20-mm diameter recording cylinder was implanted over the dorsolateral prefrontal cortex of each monkey and in the posterior parietal cortex for one monkey. Localization of the recording cylinder and electrode penetrations within the cylinder was done by registering a CT image with an MR image. Anatomical T1 images were registered to the high-resolution NMT template (NIH Macaque Template) using @animal_warper (Jung et al., 2021; Saad et al., 2009). We then used the 3D slicer software to find the projection of the recording cylinder on the brain surface. The electrode penetrations were found using the grid locations relative to the recording cylinder. Electrophysiological recordings were collected using primate Neuropixels probes with the Open Ephys data acquisition system (OpenEphys system, Atlanta, GA). For the simultaneous recordings in the prefrontal and parietal cortex, 128-contact probes from Diagnostic Biochips (Glenn Burnie, MD) were used. Electrical signals recorded from the brain were amplified, band-pass filtered between 500 and 8 kHz, and stored through a modular data acquisition system at a 30 kHz sampling rate.

## QUANTIFICATION AND STATISTICAL ANALYSIS

### Neural data processing

Recorded spike waveforms were identified and sorted into separate units using the automated spike-sorting software Kilosort (Kilosort 2.5). Single-trial peristimulus time histograms (PSTHs) for illustrations were calculated by counting spiking events of both single and multi-units in 50 ms bins. Neurons were identified as selective during any task epoch if they showed significantly different responses to the spatial location of the stimulus. This was determined using a one-way ANOVA test on the firing rates of each neuron across trials during the specific task epoch. To avoid false positives among neurons with only a few spikes, we required that a selective neuron exhibit a firing rate of at least 2 spikes per second for its best stimulus location during the task period where the ANOVA test indicated a significant main effect.

To find the preferred location of neurons, we computed the circular mean of the cue angles weighted by the neuron’s mean spike count during the delay period (Wimmer *et al*., 2014). For each neuron, we computed:

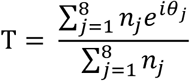

Where *n*_*j*_ is the mean spike count during the delay period in response to the cue *θ_j_* (*j*= 1…8), and we extracted its modulus *T* and angle 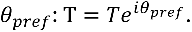 The angle *θ*_*pref*_ constitutes our estimate of the neuron’s preferred location during the delay.

### Percentage of explained variance analysis

To compute the information in a neuronal population, we used the percentage of explained variance, *ω*^2^, which measures the variance in firing rate for each neuron that can be attributed to varying the spatial location of the stimulus. The *ω*^2^statistic was calculated for every neuron in 100-ms bins with 50-ms steps, defined as follows:

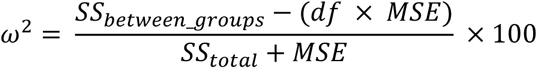

where, *SS* indicates sum of squares, MSE indicates mean squared error and *df*, degrees of freedom = 7 (we had eight stimulus conditions). To address potential bias in the calculation of PEV, we balanced the number of trials in each group by stratifying them to the lowest common number of trials across all groups.

### Laminar classification of neurons

To assess layer-specific selectivity, neurons were grouped into superficial, middle, and deep categories based on recording depth. For each penetration, the shallowest contact exhibiting spiking activity was defined as 0 mm. Depths from 0–0.8 mm were classified as superficial, the next 0.4 mm (0.8–1.2 mm) as middle, and the remaining (mostly in the range of 1.2–2.0 mm) as deep (Zhu *et al*., 2023). Within each depth bin, we quantified delay-period selectivity as the proportion of neurons that were delay-selective, computed as the number of delay-selective neurons divided by the total number of neurons recorded at the corresponding depths.

### Classification of cue location

For cross-temporal decoding, a logistic regression classifier was built with Lasso regularization (using λ = 0.01) to predict the locations of the 8 targets in the task. The firing rate matrix (estimated in 100 ms windows, overlapping by 50 ms) was z-scored across trials. Eight ‘onevsall’ classifiers were implemented by the *fitcecoc* function in Matlab, one for each of cue location. We used a ‘leave-one-out’ trial approach for decoding. Specifically, if there were *n* trials in a session, we used *n−1* trials for training at each time point and tested the model on all time points of the remaining trial. This process was repeated so that all trials were tested, and then we moved on to the next time point. Classification accuracy was defined as the proportion of correctly classified test trials.

For binary classification, we trained a binary classifier in the same way classifying each location from its opposite/diametric location. To analyze classifier confidence, we obtained the scores assigned to each target location by the classifier (using the *predict* function in Matlab), and calculated the posterior probability using:

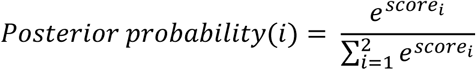

Here, i denotes the target location.

### Labeling On and Off states

To identify on and off states, we repeated the above-mentioned classification analysis for 50 times, each time randomly shuffling the training labels. For each trial, this provided us with a null distribution of confidence values. We then z-scored the true confidence values by the mean and standard deviation of the null distribution, z-values greater than 1.64 were cluster-corrected for multiple comparisons over time. Clusters with a mass greater than 95% of the null distribution were labeled as on states and contiguous z-values falling below z<0.3 threshold for 3 consecutive timepoints were labeled as Off states. For the analysis of on and off states across areas (PFC and PPC), recordings with more than 80 neurons were used. The on (or off) state was identified in one area, and then tuning and firing rate was examined in the other, for the cross-areal comparisons.

### Tuning Curves

To test how on and off states reflected in the neural activity, we labeled these states during the delay period of the task for each trial from the confidence time series and analyzed the firing rate for each unit for that particular cue location. The firing rate of each neuron, using both off- and on-states, was averaged across trials for each of the eight cue locations. Eight-element vectors for on and off-states were thus obtained for each unit. The preferred location of the unit was chosen as the cue location that had a greater firing rate after summing the on and off vectors (Panichello *et al*., 2024). The tuning functions were aligned to this location by circular rotation. Aligning tuning curves this way will naturally produce a peak at the preferred location. To correct for this, we shuffled the labels across trials 1000 times for each neuron. This yielded 1000 shuffled on and off vectors. For each shuffle, we calculated the preferred location as explained before and aligned the on and off vectors. Then we took the means of these shuffled vectors and subtracted them from the true on- and off-tuning functions. This yielded an unbiased tuning function for on and off states.

### Population Firing Rate

To illustrate how the population firing rate evolved over time, we averaged the activity of all neurons recorded from all sessions for each neuron’s preferred cue location (the preferred cue for each neuron was chosen as mentioned in the above section). In other words, we aligned each neuron’s activity across all the trials for the preferred location which yielded the mean population firing rate (Tang et al., 2019). Next, we repeated the same analysis, this time including time points labeled as On and Off.

### Inter-spike Interval Analysis

To identify coordinated silent periods within simultaneously recorded neuronal populations, we used an inter-spike interval (ISI) based method (Tao and Libedinsky, 2024). For each simultaneously recorded neuronal population, we identified delay-period selective neurons that shared the same preferred stimulus location. We then selected populations that contained at least five such neurons with overlapping selectivity, resulting in a total of n = 58 populations out of 104 possible recordings (we recorded 13 sessions, and in each session, we had 8 stimulus locations). For each population, we pooled the spike times of all neurons across all delay periods and computed the successive ISIs to generate a comprehensive empirical ISI distribution. From this, we extracted the maximum empirical ISI (MaxEmpiricalISI, size: npop × 1) for each population. In parallel, we calculated the mean firing rate (FreqEmpirical, size: npop × 1) for each population by dividing the total number of spikes by the duration of the pooled activity.

To assess whether long periods of coordinated silence exist beyond what would be expected by chance, we constructed a null distribution by shuffling neuronal activity across trials. For each of 10,000 iterations, we generated pseudo-trials by randomly mixing spike trains from different trials within a population. This procedure preserved the overall spike count and rate distribution while disrupting any trial-specific temporal coordination. From each shuffled dataset, we computed the corresponding maximum ISI value, yielding a null distribution (MaxShuffledISI, size: npop × 10,000). We also recorded the firing rate for each shuffled population (FreqShuffled, size: npop × 10,000).

To quantify the relationship between maximum ISI and firing rate, we log10-transformed both variables and performed a linear regression for each shuffled iteration, resulting in a distribution of 10,000 slopes and intercepts. We then performed a comparable regression using the empirical data (log10(MaxEmpiricalISI) vs. log10(FreqEmpirical)) to obtain the empirical slope and intercept. To validate our approach and assess its sensitivity to coordinated silent periods, we generated surrogate neural population data with and without imposed silent periods. Each simulated population consisted of a randomly chosen number of neurons (uniformly sampled from 2 to 20) and trials (uniformly sampled from 10 to 20). Each trial was 3s long same as in our task. We simulated a total of 50 such populations.

For each simulated neuron, a baseline firing rate was randomly drawn from a discrete uniform distribution ranging from 1 to 20 Hz. This firing rate determined the expected number of spikes per second for generating spike trains using a Poisson process. For each trial, we optionally inserted silent periods—contiguous windows of suppressed activity—by removing spikes within these intervals. Each trial was assigned a random number of silent periods (between 1 and 3), with each period lasting 100 ms. The onset times of silent periods were drawn uniformly from the trial duration to ensure random placement. For each population, we produced two versions of spike data: without off-(silence) periods, created by Poisson spike trains without modification; and with silence, created by Poisson spike trains from which spikes were removed within the silent windows. The resulting data structures contained spike times for each neuron across all trials, stored separately for the two conditions (with and without silent periods). These surrogate datasets were then analyzed using the same population ISI procedure described above, including computation of max ISI values and construction of null distributions through 10,000 trial-shuffled surrogates per population. This simulation allowed us to test whether our method could reliably distinguish datasets with coordinated silent periods from those without.

### Mixture modeling of confidence

We tested whether confidence during the delay period was better captured by a one-state versus two-state model using mixture modeling. For each session and cue location, we fit (i) a single beta distribution:

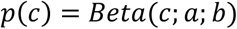

and (ii) a two-component beta mixture:

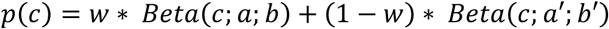

to the distribution of confidence values in the delay. Model comparison was based on the Akaike Information Criterion (AIC) and Bayesian Information Criterion (BIC): we computed ΔAIC = AIC2-state − AIC1-state and ΔBIC = BIC2-state − BIC1-state, with negative ΔAIC and ΔBIC values indicating better fit for 2-state model.

### Power spectrum of on/off states

To assess whether the on/off states exhibited rhythmicity, we generated, for each trial and state (“off” or “on”), a binary time series marking time points belonging to that state and mean-centered each series. To extend the frequency range and capture broadband rhythms, we repeated the binary classification using 10-ms bins (sampling rate 100 Hz), enabling spectral estimates up to 50 Hz. We then computed one-sided power spectral densities (PSDs) using Welch’s method with 2.0 s Hamming windows with 50% overlap and a 1024-point FFT (zero-padding for frequency resolution). PSDs were estimated separately for PFC and PPC and for each state on a trial-by-trial basis and summarized by averaging the PSD across trials at each frequency.

### Reaction time vs state analysis

To investigate the relationship between reaction time and the on/off state, for each session, we computed reaction time (RT; in ms) and marked whether the neural state at fixation offset (the “go” cue for the saccade) was “on” (State=1) or “off” (State=0). Trials with missing state labels were excluded. Within each session, we fit a generalized linear model to quantify the effect of state on RT. The session-level coefficient for State (β_state) represents the difference in mean RT between “on” and “off” states (RT_on − RT_off). To obtain population-level inference while accounting for session-level clustering, we performed a nonparametric bootstrap over sessions (B = 10,000 draws). For each bootstrap draw, we sampled sessions with replacement and computed the mean of the corresponding β_state values. This yielded a bootstrap distribution of the mean state effect. To assess whether the mean effect was positive, we computed one-sided p-values as the proportion of bootstrap means ≥ 0. As defined, a positive β_state indicates longer RTs when the state is “on” relative to “off”; a negative β_state indicates shorter RTs in the “on” state.

### On and off states in correct and error trials

We calculated the number of on and off states per trial in correct and error trials. For each session, we calculated the site’s overall preferred cue location (that produced the greatest mean classification confidence during the delay period) and the least-preferred cue condition (smallest mean classification confidence during the delay period). Then we counted the number of on and off states that occurred in the last 0.5s of the delay period per trial in these two conditions. To obtain population-level inference, we again adopted a bootstrap method (as explained in the previous section). Thus, it yielded a distribution of mean states per trial.

## ACKNOWLEDGEMENTS

Supported by NIH award number R01 EY017077 and NSF award 2011514. We wish to thank Chrissy Suell and Kayla Yetman for technical help and Albert Compte, Melanie Tschiersch, Ethan Meyers, Tirin Moore, and Matt Panichello for helpful comments on an earlier version of this manuscript.

## AUTHOR CONTRIBUTIONS

CC designed the experiment. RM, WD, JZ, BJ, and AM performed behavioral training and neurophysiological recordings. RM, and ZW performed data analyses. CC and RM wrote the paper with input from all authors.

## DECLARATION OF GENERATIVE AI AND AI-ASSISTED TECHNOLOGIES

No AI tools were used during the preparation of this work

## SUPPLEMENTAL INFORMATION

Document S1. Figures S1–S10

## SUPPLEMENTAL FIGURES

**Figure S1.**
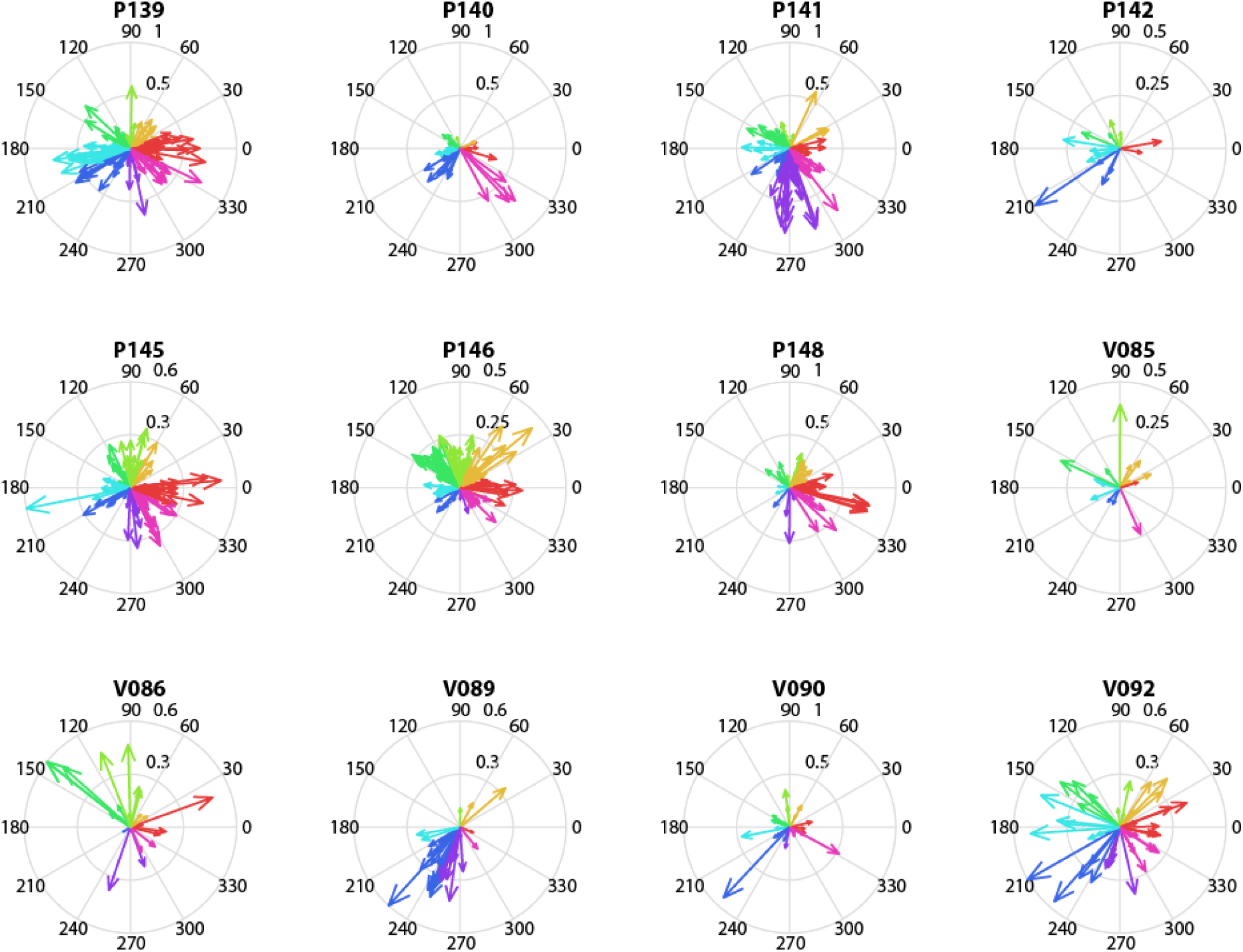
Population compass plots, related to Figure 1. Plots illustrate the tuning of all delay-selective neurons across all recording sessions analyzed (in addition to session P138, shown in Fig. 1).

**Figure S2.**
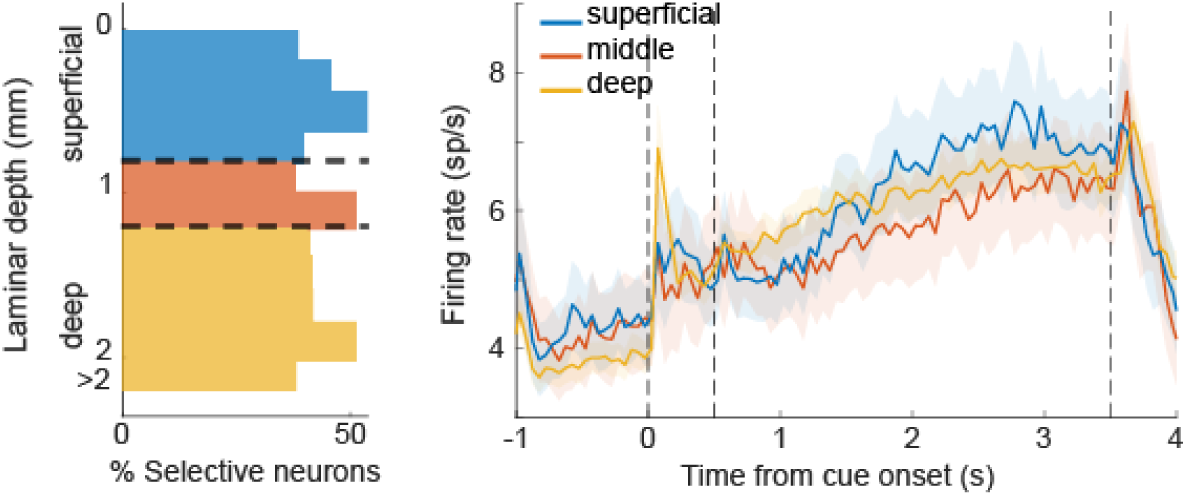
Laminar distribution of delay activity, related to Figure 1. (A). Histogram represents the percentage of selective neurons recorded at each depth relative to the top of the cortex (depth 0) that exhibited tuned, persistent activity. B. Population PSTH for neurons divided in superficial (0-800 μm, n=149), middle (800-1200 μm, n=119) and deep layers (>1200 μm from the top of the cortex, n=667).

**Figure S3.**
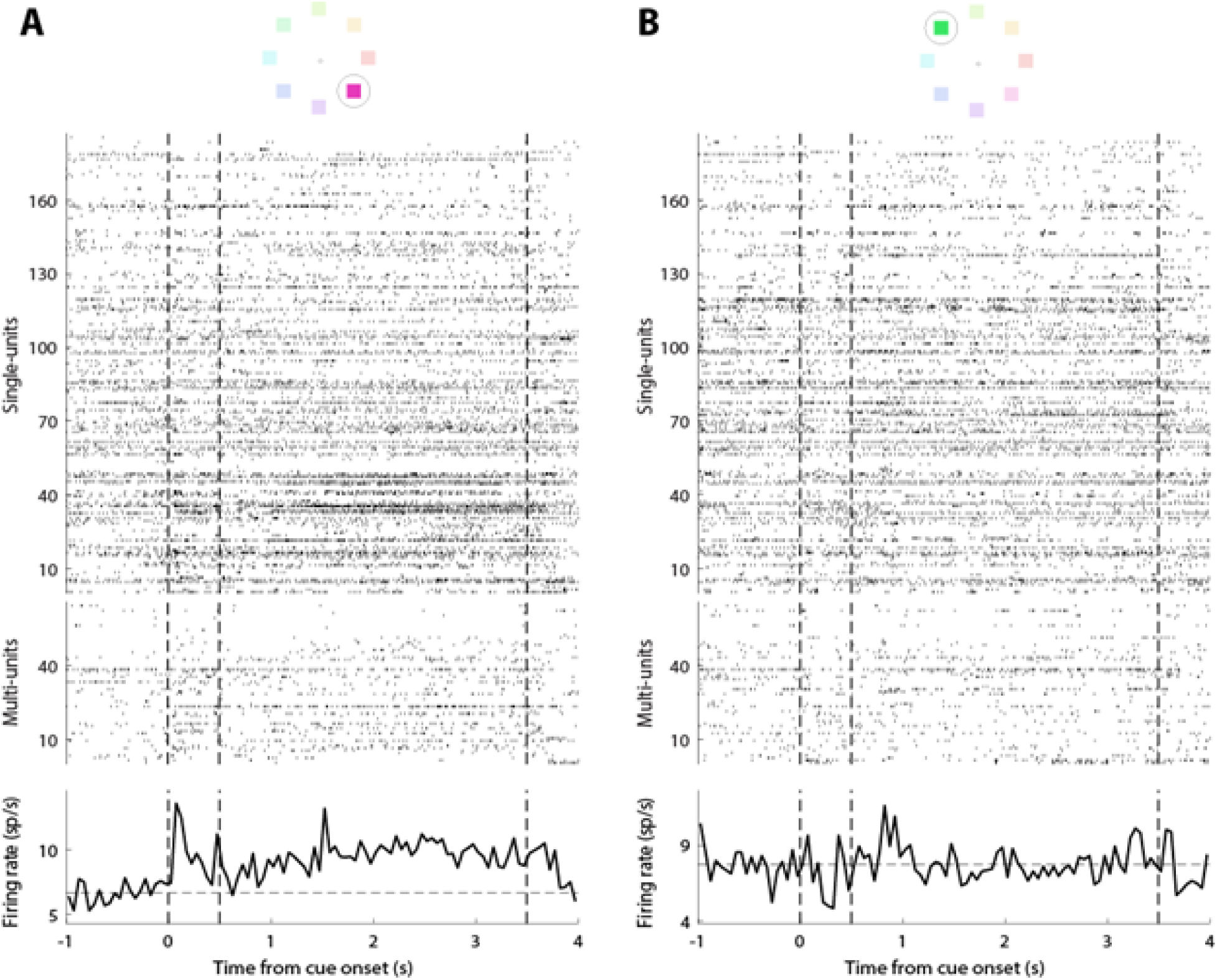
Examples of persistent and non-persistent delay activity, related to Figure 1. Raster plot and PSTH for two example trials (left: 315° cue stimulus, right: 135° cue stimulus) for single and multi-units recorded simultaneously from a recording session (P138). The conventions are the same as in Figure 2C.

**Figure S4.**
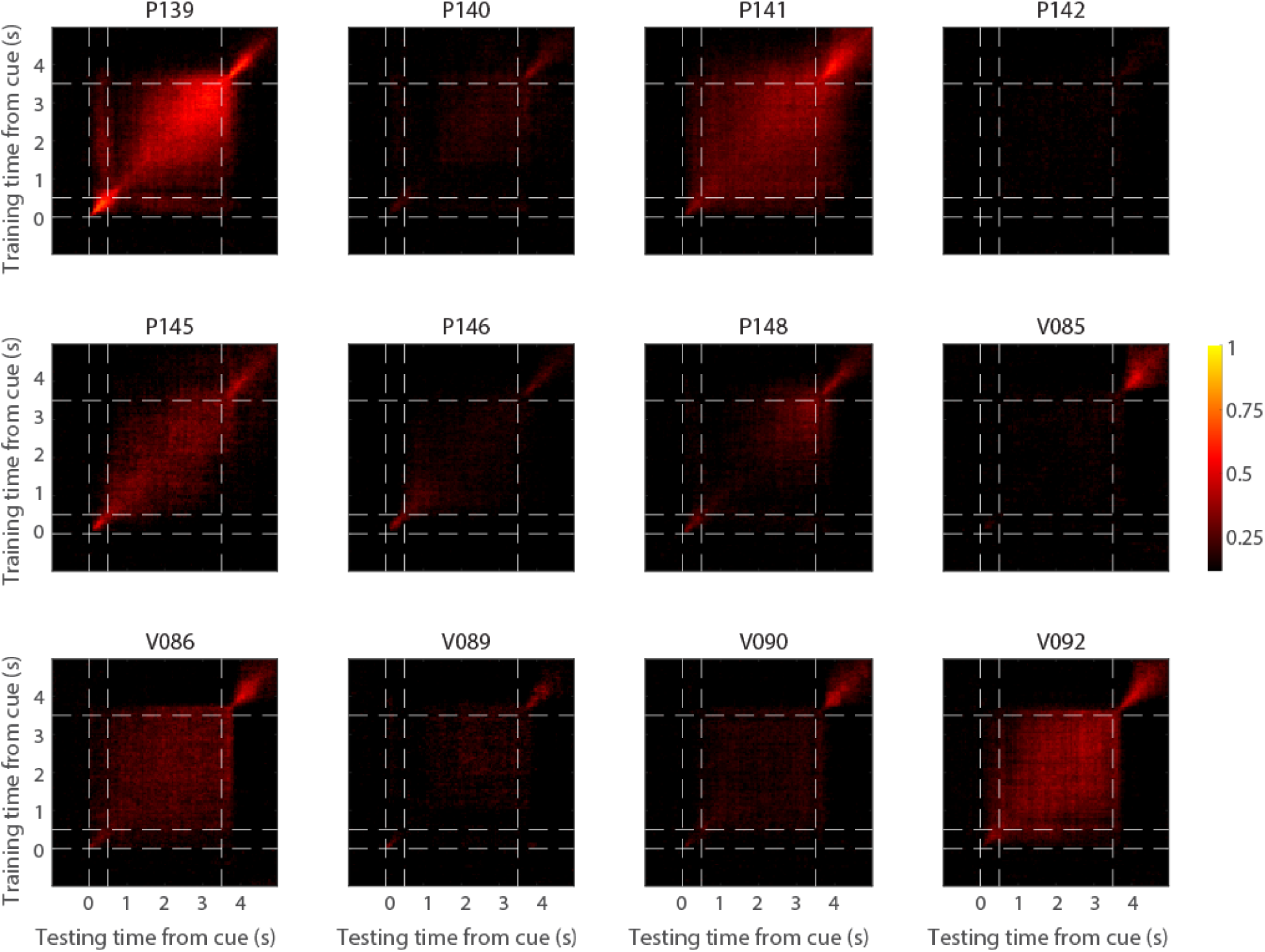
Cross-temporal decoding, related to Figure 2. Mean cross-temporal decoding for all recording sessions. The conventions are the same as in Figure 2 (central panel).

**Figure S5.**
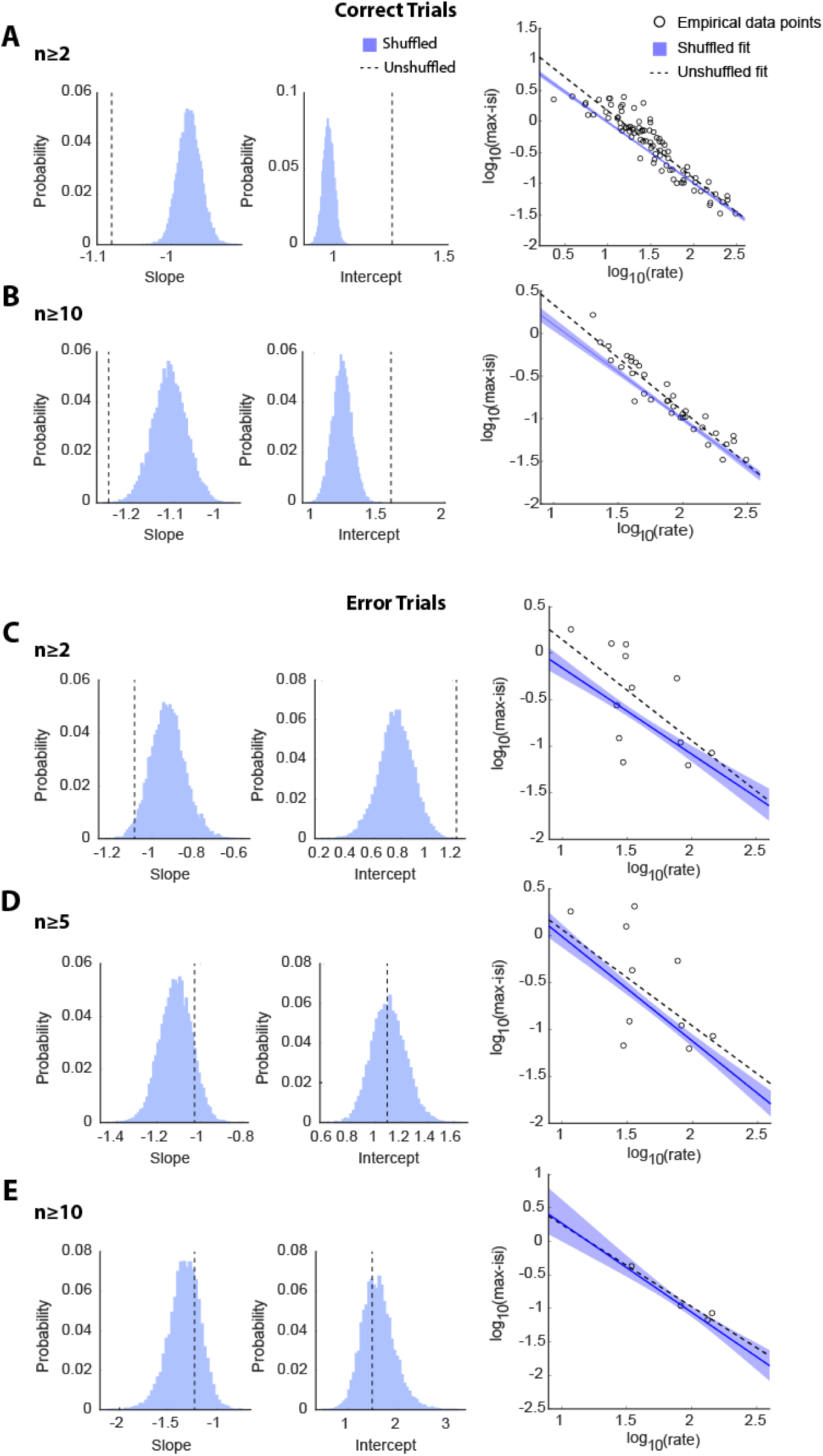
ISI analysis in populations of varied size and in error trials, related to Figure 3. (A-B) Same as in Fig. 3I-K, for ISIs computed from populations of at least 2 (A) or 10 neurons (B). (C-E). Same as above, but for error trials of populations with at least 2 (C), 5 (D), and 10 (E) neurons.

**Figure S6.**
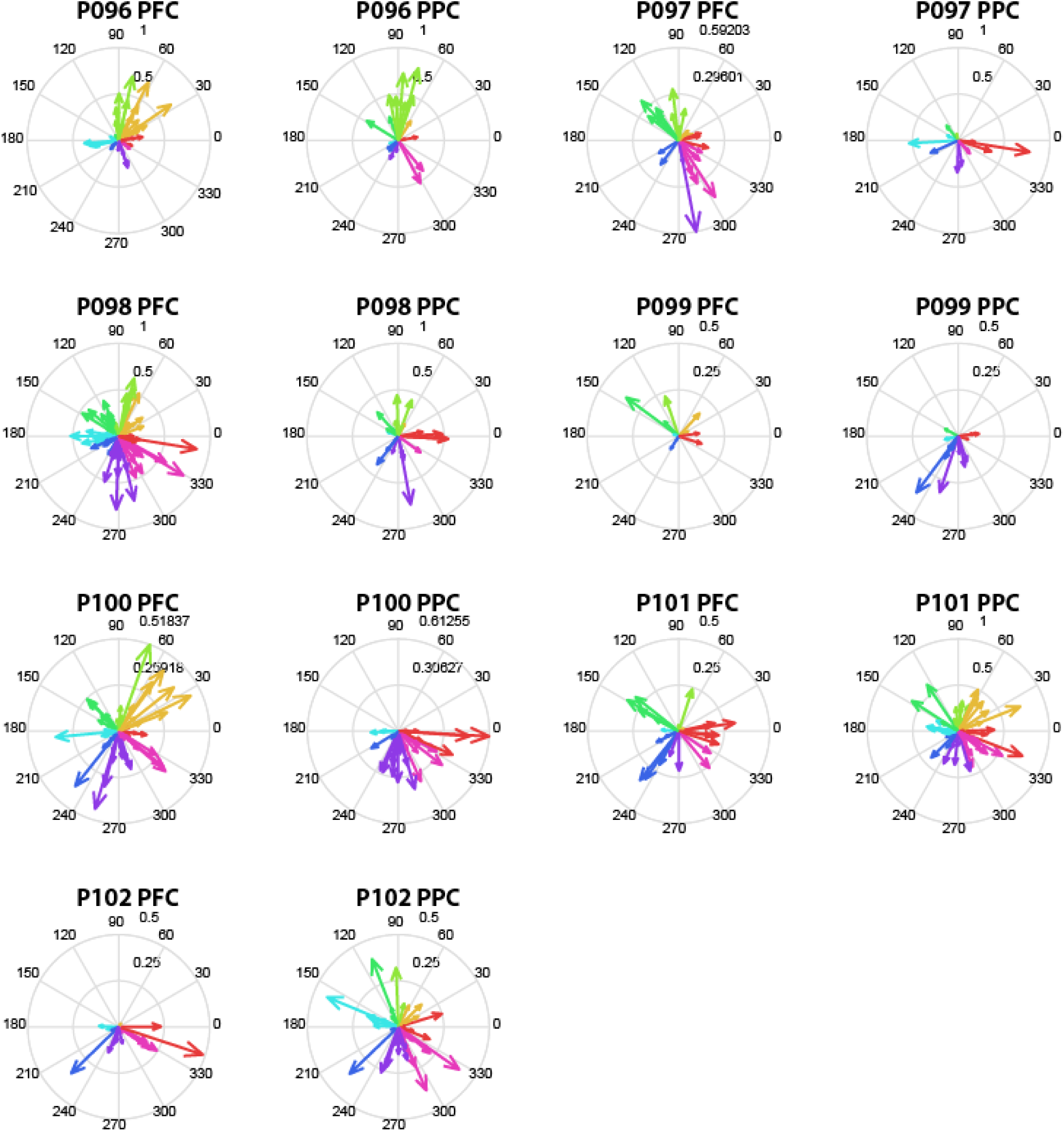
Compass plots for simultaneous PFC-PPC recordings, related to Figure 4. Compass plots are shown for all neurons recorded simultaneously in sessions with PFC-PPC recordings.

**Figure S7.**
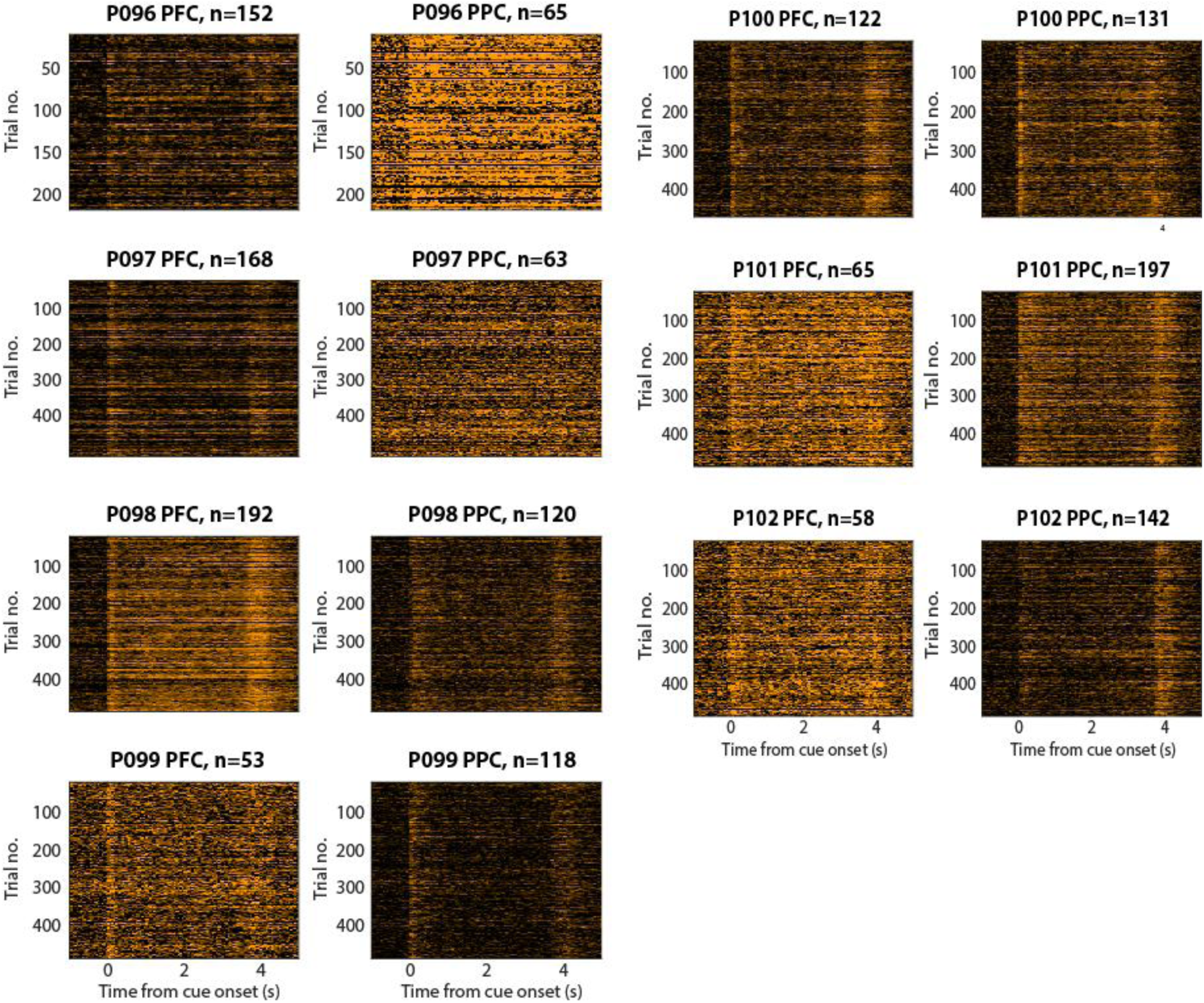
Single-trial classifier confidence for simultaneous PFC-PPC recordings, related to Figure 4. Single-trial classifier confidence is shown. Color scale as in Fig. 4A.

**Figure S8.**
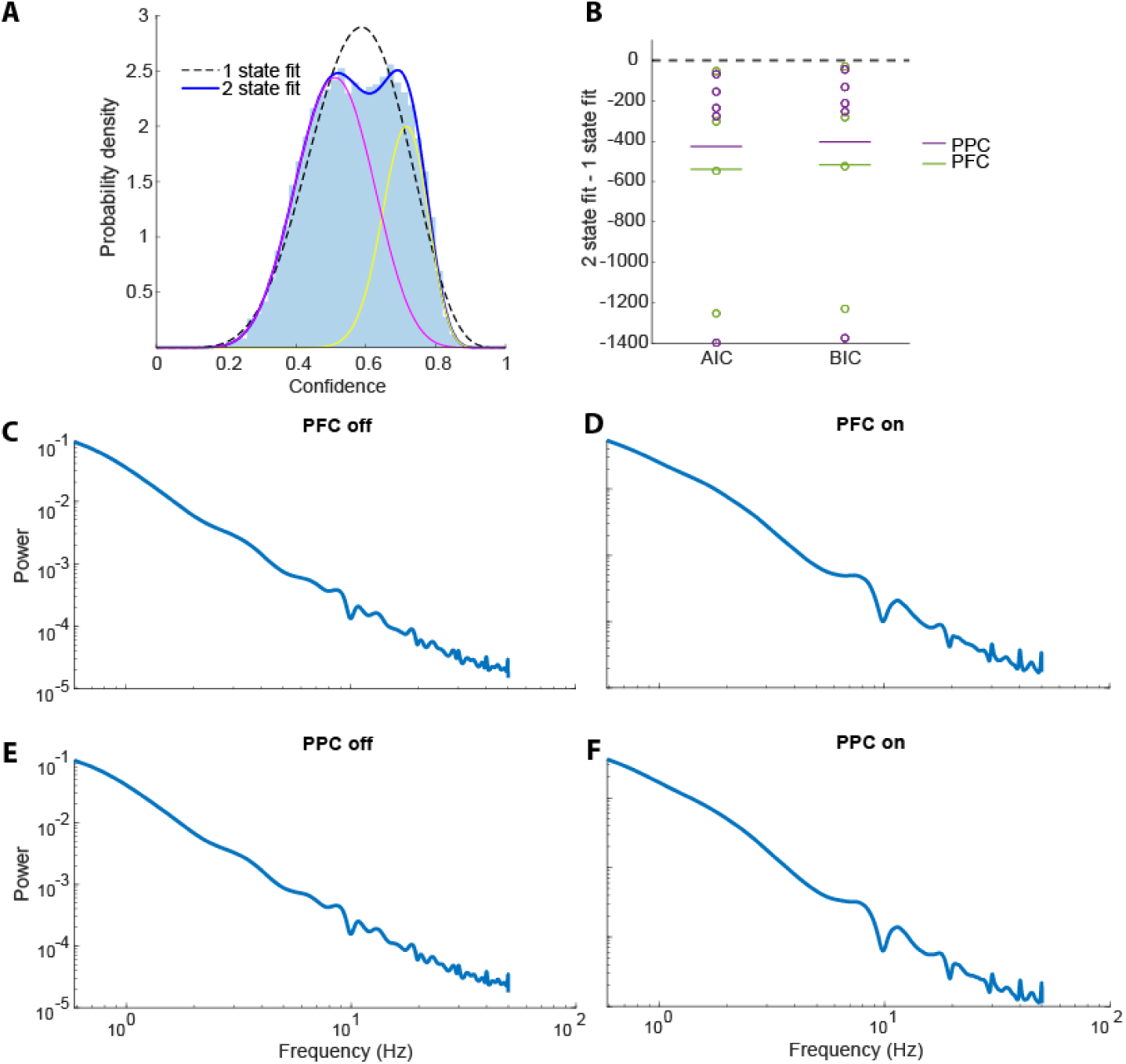
Characteristics of on- and off-states, related to Figure 4. (A). Example histogram of confidence values for one cue location from an example PFC session. The dashed line shows the 1-state beta fit while the solid blue lines demonstrate a 2-state fit. The magenta and yellow lines show the components for the two-state model. (B). Akaike Information criterion (AIC) and Bayesian Information Criterion (BIC) values for 1- and 2-state fits for all PPC and PFC sessions. (C-F). Welch power spectral density is plotted for on- and off-states in the prefrontal and posterior parietal cortex.

**Figure S9.**
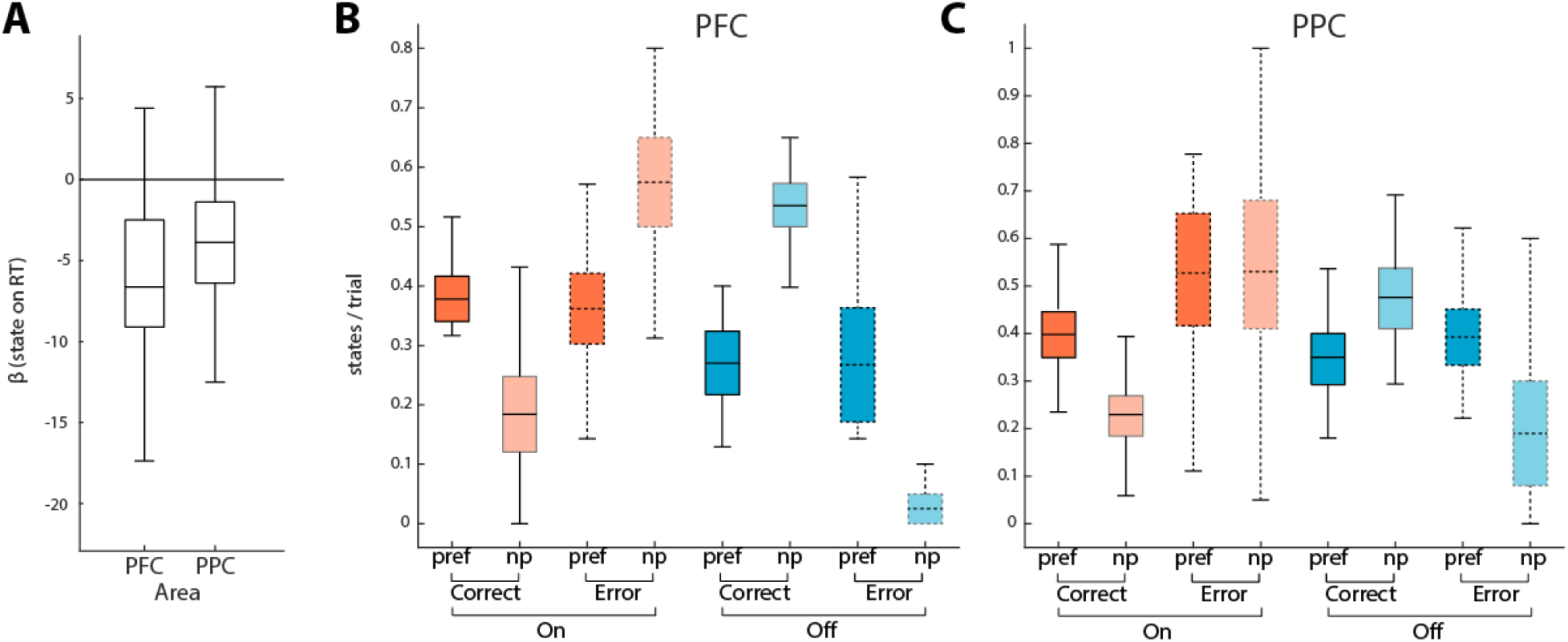
Relationship of on- and off-states with behavior, related to Figure 4. (A). Regression coefficient values for on- and off-states with relation to reaction time (RT). Negative values indicate shorter reaction times during on-states. (B). Number of on- (red) and off-states (blue) in the PFC per trial, for correct (solid boxes) and error trials (dotted boxes), when the stimulus appeared at the overall preferred location of the site (pref - darker colors) or non-preferred location (np - lighter colors). (C). Same as B for the PPC.

**Figure S10.**
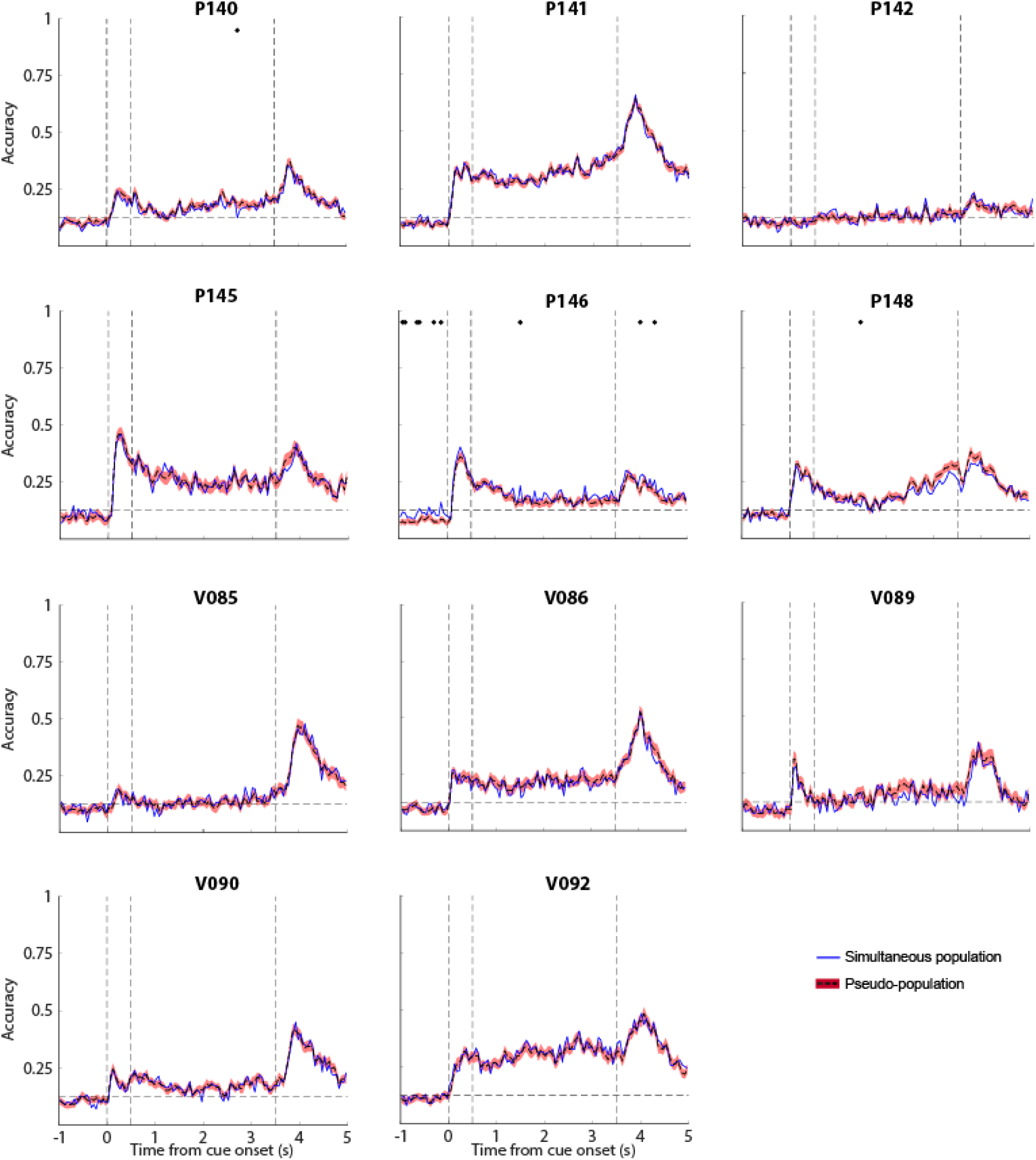
Decoding of simultaneous and pseudo-populations, related to Figure 5. Mean decoding performance is plotted for PFC recording sessions for populations of neurons obtained simultaneously vs. pseudo-populations constructed by splicing different trials together. Conventions are the same as in Fig. 5.

## REFERENCES

Baddeley, A. (2012). Working memory: theories, models, and controversies. Annu. Rev. Psychol. 63, 1–29. 10.1146/annurev-psych-120710-100422.

Barbosa, J., Stein, H., Martinez, R.L., Galan-Gadea, A., Li, S., Dalmau, J., Adam, K.C.S., Valls-Sole, J., Constantinidis, C., and Compte, A. (2020). Interplay between persistent activity and activity-silent dynamics in the prefrontal cortex underlies serial biases in working memory. Nat. Neurosci. 23, 1016–1024. 10.1038/s41593-020-0644-4.

Brunel, N. (2000). Dynamics of sparsely connected networks of excitatory and inhibitory spiking neurons. J. Comput. Neurosci. 8, 183–208.

Chaudhuri, R., and Fiete, I. (2016). Computational principles of memory. Nat. Neurosci. 19, 394–403. 10.1038/nn.4237.

Cohen, M.R., and Kohn, A. (2011). Measuring and interpreting neuronal correlations. Nat. Neurosci. 14, 811–819. nn.2842

Compte, A., Brunel, N., Goldman-Rakic, P.S., and Wang, X.J. (2000). Synaptic mechanisms and network dynamics underlying spatial working memory in a cortical network model. Cereb. Cortex 10, 910–923.

Compte, A., Constantinidis, C., Tegner, J., Raghavachari, S., Chafee, M.V., Goldman-Rakic, P.S., and Wang, X.J. (2003). Temporally irregular mnemonic persistent activity in prefrontal neurons of monkeys during a delayed response task. J. Neurophysiol. 28, 3441–3454.

Constantinidis, C., Franowicz, M.N., and Goldman-Rakic, P.S. (2001a). Coding specificity in cortical microcircuits: a multiple electrode analysis of primate prefrontal cortex. J. Neurosci. 21, 3646–3655.

Constantinidis, C., Franowicz, M.N., and Goldman-Rakic, P.S. (2001b). The sensory nature of mnemonic representation in the primate prefrontal cortex. Nat. Neurosci. 4, 311–316.

Constantinidis, C., Funahashi, S., Lee, D., Murray, J.D., Qi, X.L., Wang, M., and Arnsten, A.F.T. (2018). Persistent Spiking Activity Underlies Working Memory. J. Neurosci. 38, 7020–7028. 10.1523/JNEUROSCI.2486-17.2018.

Constantinidis, C., and Goldman-Rakic, P.S. (2002). Correlated discharges among putative pyramidal neurons and interneurons in the primate prefrontal cortex. J. Neurophysiol. 88, 3487–3497.

Davis, Z.W., Muller, L., Martinez-Trujillo, J., Sejnowski, T., and Reynolds, J.H. (2020). Spontaneous travelling cortical waves gate perception in behaving primates. Nature 587, 432–436. 10.1038/s41586-020-2802-y.

Ebitz, R.B., and Hayden, B.Y. (2021). The population doctrine in cognitive neuroscience. Neuron. 10.1016/j.neuron.2021.07.011.

Funahashi, S., Bruce, C.J., and Goldman-Rakic, P.S. (1989). Mnemonic coding of visual space in the monkey’s dorsolateral prefrontal cortex. J. Neurophysiol. 61, 331–349.

Fuster, J.M., and Alexander, G.E. (1971). Neuron activity related to short-term memory. Science 173, 652–654.

Hansel, D., and Mato, G. (2003). Asynchronous states and the emergence of synchrony in large networks of interacting excitatory and inhibitory neurons. Neural Comput. 15, 1–56.

Hansel, D., and Sompolinsky, H. (1996). Chaos and synchrony in a model of a hypercolumn in visual cortex. J. Comput. Neurosci. 3, 7–34.

Jaffe, R.J., and Constantinidis, C. (2021). Working Memory: From Neural Activity to the Sentient Mind. Compr Physiol 11, 1–41. 10.1002/cphy.c210005.

Jun, J.J., Steinmetz, N.A., Siegle, J.H., Denman, D.J., Bauza, M., Barbarits, B., Lee, A.K., Anastassiou, C.A., Andrei, A., Aydin, C., et al. (2017). Fully integrated silicon probes for high-density recording of neural activity. Nature 551, 232–236. 10.1038/nature24636.

Jung, B., Taylor, P.A., Seidlitz, J., Sponheim, C., Perkins, P., Ungerleider, L.G., Glen, D., and Messinger, A. (2021). A comprehensive macaque fMRI pipeline and hierarchical atlas. NeuroImage 235, 117997. 10.1016/j.neuroimage.2021.117997.

Lansner, A., Fiebig, F., and Herman, P. (2023). Fast Hebbian plasticity and working memory. Curr. Opin. Neurobiol. 83, 102809. 10.1016/j.conb.2023.102809.

Lara, A.H., and Wallis, J.D. (2014). Executive control processes underlying multi-item working memory. Nat. Neurosci. 17, 876–883. 10.1038/nn.3702.

Leavitt, M.L., Pieper, F., Sachs, A.J., and Martinez-Trujillo, J.C. (2017). Correlated variability modifies working memory fidelity in primate prefrontal neuronal ensembles. Proc. Natl. Acad. Sci. U. S. A. 114, E2494–E2503. 10.1073/pnas.1619949114.

Lundqvist, M., Compte, A., and Lansner, A. (2010). Bistable, irregular firing and population oscillations in a modular attractor memory network. PLoS Comput. Biol. 6, e1000803. 10.1371/journal.pcbi.1000803.

Lundqvist, M., Herman, P., and Miller, E.K. (2018). Working Memory: Delay Activity, Yes! Persistent Activity? Maybe Not. J. Neurosci. 38, 7013–7019. 10.1523/JNEUROSCI.2485-17.2018.

Lundqvist, M., Rose, J., Herman, P., Brincat, S.L., Buschman, T.J., and Miller, E.K. (2016). Gamma and Beta Bursts Underlie Working Memory. Neuron 90, 152–164. 10.1016/j.neuron.2016.02.028.

Markowitz, D.A., Curtis, C.E., and Pesaran, B. (2015). Multiple component networks support working memory in prefrontal cortex. Proc. Natl. Acad. Sci. U. S. A. 112, 11084–11089. 10.1073/pnas.1504172112.

Mejias, J.F., and Wang, X.J. (2022). Mechanisms of distributed working memory in a large-scale network of macaque neocortex. Elife 11, 72136. 10.7554/eLife.72136.

Mendoza-Halliday, D., and Martinez-Trujillo, J.C. (2017). Neuronal population coding of perceived and memorized visual features in the lateral prefrontal cortex. Nat Commun 8, 15471. 10.1038/ncomms15471.

Mendoza-Halliday, D., Torres, S., and Martinez-Trujillo, J.C. (2014). Sharp emergence of feature-selective sustained activity along the dorsal visual pathway. Nat. Neurosci. 17, 1255–1262. 10.1038/nn.3785.

Meyer, T., and Constantinidis, C. (2005). A software solution for the control of visual behavioral experimentation. Journal of Neuroscience Methods 142, 27–34. 10.1016/j.jneumeth.2004.07.009.

Meyers, E.M., Qi, X.L., and Constantinidis, C. (2012). Incorporation of new information into prefrontal cortical activity after learning working memory tasks. Proc. Natl. Acad. Sci. U. S. A. 109, 4651–4656. 1201022109.

Miller, E.K., Lundqvist, M., and Bastos, A.M. (2018). Working Memory 2.0. Neuron 100, 463–475. 10.1016/j.neuron.2018.09.023.

Mongillo, G., Barak, O., and Tsodyks, M. (2008). Synaptic theory of working memory. Science 319, 1543–1546.

Nassar, M.R., Rumsey, K.M., Wilson, R.C., Parikh, K., Heasly, B., and Gold, J.I. (2012). Rational regulation of learning dynamics by pupil-linked arousal systems. Nat. Neurosci. 15, 1040–1046. 10.1038/nn.3130.

Ostojic, S. (2014). Two types of asynchronous activity in networks of excitatory and inhibitory spiking neurons. Nat. Neurosci. 17, 594–600. 10.1038/nn.3658.

Panichello, M.F., Jonikaitis, D., Oh, Y.J., Zhu, S., Trepka, E.B., and Moore, T. (2024). Intermittent rate coding and cue-specific ensembles support working memory. Nature 636, 422–429. 10.1038/s41586-024-08139-9.

Pesaran, B., Pezaris, J.S., Sahani, M., Mitra, P.P., and Andersen, R.A. (2002). Temporal structure in neuronal activity during working memory in macaque parietal cortex. Nat. Neurosci. 5, 805–811.

Qi, X.L., and Constantinidis, C. (2012). Correlated discharges in the primate prefrontal cortex before and after working memory training Eur. J. Neurosci. 36, 3538–3548.

Renart, A., de la Rocha, J., Bartho, P., Hollender, L., Parga, N., Reyes, A., and Harris, K.D. (2010). The asynchronous state in cortical circuits. Science 327, 587–590. 327/5965/587.

Rose, N.S., LaRocque, J.J., Riggall, A.C., Gosseries, O., Starrett, M.J., Meyering, E.E., and Postle, B.R. (2016). Reactivation of latent working memories with transcranial magnetic stimulation. Science 354, 1136–1139. 10.1126/science.aah7011.

Saad, Z.S., Glen, D.R., Chen, G., Beauchamp, M.S., Desai, R., and Cox, R.W. (2009). A New Method for Improving Functional-to-Structural MRI Alignment using Local Pearson Correlation. NeuroImage 44, 839–848. 10.1016/j.neuroimage.2008.09.037.

Stokes, M.G. (2015). ’Activity-silent’ working memory in prefrontal cortex: a dynamic coding framework. Trends Cogn. Sci. 19, 394–405. 10.1016/j.tics.2015.05.004.

Stokes, M.G., Kusunoki, M., Sigala, N., Nili, H., Gaffan, D., and Duncan, J. (2013). Dynamic coding for cognitive control in prefrontal cortex. Neuron 78, 364–375. 10.1016/j.neuron.2013.01.039.

Sugase-Miyamoto, Y., Liu, Z., Wiener, M.C., Optican, L.M., and Richmond, B.J. (2008). Short-term memory trace in rapidly adapting synapses of inferior temporal cortex. PLoS Comput. Biol. 4, e1000073. 10.1371/journal.pcbi.1000073.

Tan, A.Y., Chen, Y., Scholl, B., Seidemann, E., and Priebe, N.J. (2014). Sensory stimulation shifts visual cortex from synchronous to asynchronous states. Nature 509, 226–229. 10.1038/nature13159.

Tang, H., Qi, X.L., Riley, M.R., and Constantinidis, C. (2019). Working memory capacity is enhanced by distributed prefrontal activation and invariant temporal dynamics. Proc. Natl. Acad. Sci. U. S. A. 116, 7095–7100. 10.1073/pnas.1817278116.

Tao, W., and Libedinsky, C. (2024). Evidence of Activity-Silent Working Memory in Prefrontal Cortex. bioRxiv 10.1101/2024.06.03.597259.

Wang, X.J. (2021). 50 years of mnemonic persistent activity: quo vadis? Trends Neurosci. 44, 888–902. 10.1016/j.tins.2021.09.001.

Watanabe, K., and Funahashi, S. (2007). Prefrontal delay-period activity reflects the decision process of a saccade direction during a free-choice ODR task. Cerebral Cortex 17, i88–i100.

Wimmer, K., Nykamp, D.Q., Constantinidis, C., and Compte, A. (2014). Bump attractor dynamics in prefrontal cortex explains behavioral precision in spatial working memory. Nat. Neurosci. 17, 431–439. 10.1038/nn.3645.

Wolff, M.J., Jochim, J., Akyürek, E.G., and Stokes, M.G. (2017). Dynamic hidden states underlying working-memory-guided behavior. Nature neuroscience 20, 864–871.

Zaksas, D., and Pasternak, T. (2006). Directional signals in the prefrontal cortex and in area MT during a working memory for visual motion task. J. Neurosci. 26, 11726–11742. 26/45/11726.

Zanos, T.P., Mineault, P.J., Nasiotis, K.T., Guitton, D., and Pack, C.C. (2015). A sensorimotor role for traveling waves in primate visual cortex. Neuron 85, 615–627. 10.1016/j.neuron.2014.12.043.

Zhou, X., Katsuki, F., Qi, X.L., and Constantinidis, C. (2012). Neurons with inverted tuning during the delay periods of working memory tasks in the dorsal prefrontal and posterior parietal cortex. J. Neurophysiol. 108, 31–38. jn.01151.2011.

Zhou, X., Zhu, D., Qi, X.L., Lees, C.J., Bennett, A.J., Salinas, E., Stanford, T.R., and Constantinidis, C. (2013). Working Memory Performance and Neural Activity in the Prefrontal Cortex of Peri-pubertal Monkeys. J. Neurophysiol. 110, 2648–2660. doi: 10.1152/jn.00370.2013.

Zhu, J., Hammond, B.M., Zhou, X.M., and Constantinidis, C. (2023). Laminar pattern of adolescent development changes in working memory neuronal activity. J. Neurophysiol. 130, 980–989. 10.1152/jn.00294.2023.

Zohary, E., Shadlen, M.N., and Newsome, W.T. (1994). Correlated neuronal discharge rate and its implications for psychophysical performance. Nature 370, 140–143.

Zylberberg, J., and Strowbridge, B.W. (2017). Mechanisms of Persistent Activity in Cortical Circuits: Possible Neural Substrates for Working Memory. Annu. Rev. Neurosci. 40, 603–627. 10.1146/annurev-neuro-070815-014006.

